# Intra-host IRES heterogeneity shapes hepatitis C virus translation through epistatic and population-level effects

**DOI:** 10.64898/2026.08.05.743084

**Authors:** Natalia Echeverría, Paula Perbolianachis, Fabiana Gámbaro, Lucas Amaya, Martín Soñora, Alvaro Fajardo, Nelia Hernández, Juan Cristina, Evandro Ferrada, Pilar Moreno, Gonzalo Moratorio

**Affiliations:** Laboratorio de Virología Molecular, Centro de Investigaciones Nucleares, Facultad de Ciencias, Universidad de la República. Montevideo, Uruguay; Laboratorio de Evolución Experimental de Virus, Institut Pasteur de Montevideo. Montevideo, Uruguay; Clínica de Gastroenterología, Hospital de Clínicas, Facultad de Medicina, Universidad de la República. Montevideo, Uruguay; Instituto de Neurociencia, Facultad de Ciencias, Universidad de Valparaíso, Valparaíso, Chile; Centro Interdisciplinario de Neurociencia de Valparaíso, Facultad de Ciencias, Universidad de Valparaíso, Valparaíso, Chile; Instituto de Sistemas Complejos de Valparaíso (ISCV), Artillería 470, Cerro Artillería, Valparaíso, Chile

## Abstract

Hepatitis C virus (HCV) circulates within each patient as a diverse population of closely related genomes, yet RNA functional properties are commonly inferred from a single consensus or dominant genome. The contribution of non-coding intra-host variability, particularly within the internal ribosome entry site (IRES), to translational efficiency remains poorly defined. Here we investigated how naturally occurring HCV IRES variation influences viral RNA translation. Complete IRES sequences from chronically infected patients were analyzed using molecular cloning, bicistronic reporters, full-length replication-deficient viral RNAs and reconstructed intra-host populations. We found that natural IRES mutations displayed context-dependent effects, and combinations of mutations produced translational phenotypes that could not be predicted from the corresponding single mutations, consistent with intragenic epistasis. Moreover, several variants behaved differently in bicistronic reporters and full-length viral RNAs, demonstrating that both genomic and cellular context shape IRES function. Reconstructed genotype 1a populations largely reproduced the activity of their dominant haplotypes. In contrast, reconstructed genotype 3a populations translated substantially more efficiently than their corresponding dominant sequences, showing that low-frequency variants can collectively modulate translation at the population level. These findings demonstrate that the translational phenotype of HCV cannot always be inferred from the dominant sequence alone and identify epistasis, genomic context, and intra-host population composition as interacting determinants of viral RNA translation.

**Importance:** Hepatitis C virus (HCV) exists within each infected person as a diverse population of closely related viruses rather than as a single genetic sequence. This study shows that natural variation in a key RNA region controlling viral protein production can alter how efficiently the virus functions, and that these effects depend on combinations of mutations rather than on individual changes alone. By analyzing complete viral RNAs in addition to widely used reporter systems, we demonstrate that the full viral genome can substantially influence the activity of this regulatory region, providing a more realistic view of how translation occurs during natural infection. Our findings also reveal that rare viral variants can collectively shape the behavior of the viral population, challenging the common practice of relying on a single dominant sequence to represent an infection. These results provide new insight into how genetic diversity drives HCV evolution and adaptation.

## Introduction

Hepatitis C virus (HCV) is a positive-sense RNA virus of the family *Flaviviridae* that establishes chronic infection and remains a major cause of liver disease worldwide(1, 2). HCV populations display extensive genetic diversity both among infected individuals and within each host, with consequences for viral evolution, clinical outcome, and treatment response(3). Although direct-acting antivirals achieve cure rates above 95% with improved safety(4), limited access to diagnosis and therapy and the absence of a vaccine remain major barriers to HCV control(5, 6).

Translation is the first biosynthetic event after the HCV genome enters the cytoplasm and is initiated by the internal ribosome entry site (IRES)(7). HCV IRES is a highly structured RNA element located within the 5’ untranslated region (UTR) and extending into the core coding sequence. The IRES structure enables direct recruitment of the ribosome to the viral RNA(8), bypassing the requirement for a 5’ cap. This mechanism drives translation of a single polyprotein, which is subsequently cleaved into 10 structural and non-structural proteins required for virion assembly and viral replication, respectively(9).

HCV IRES-mediated translation is determined by both intrinsic RNA features and the cellular environment. Its activity is modulated by multiple factors, including IRES secondary structure(10–12), RNA sequence variation(13), RNA modifications(14), long-range interactions(15–18), and IRES trans-acting factors (ITAFs), such as La(19, 20), PTB1/4(21), nucleolin(22), NSAP1(23, 24), hnRNP L/D(25, 26), Gemin 5(27) and PSPC1(28).

In addition, miR-122 indirectly enhances translation by stabilizing functional IRES conformations(29). Understanding how these factors shape translation is central to defining HCV replication strategies and potential antiviral vulnerabilities(30, 31).

Most functional studies of HCV IRES variation have examined consensus sequences or individual cloned variants. However, HCV, like other RNA viruses, circulates as a genetically diverse intra-host population in which a dominant haplotype coexists with numerous low-frequency variants that contribute to adaptation and can influence viral fitness, replication, and cellular tropism(32–37). Consequently, representing these populations by a consensus sequence or the most abundant haplotype may be particularly limiting for structured noncoding RNA elements, where the effects of nucleotide substitutions depend on sequence context and low-frequency variants may contribute to the phenotype of the viral population as a whole.

Previous studies have reported differences in translational activity among naturally occurring HCV IRES sequences and among variants derived from distinct tissues or cellular environments(38–42). Moreover, mixtures of patient-derived IRES variants display translational properties distinct from those of individual clones, suggesting that low-frequency variants collectively influence population-level translational output(43). Consistent with this concept, group selection has been described in RNA viruses, highlighting contributions of minority genomes to overall population fitness(44).

However, all studies investigating HCV IRES-mediated translation relied exclusively on bicistronic reporter constructs, leaving unresolved whether naturally occurring IRES mutations interact epistatically and whether these population-level effects are preserved in more physiologically relevant viral RNA contexts. A deeper understanding of these mechanisms is essential for elucidating the molecular basis of HCV infection and persistence.

Here, we combined experimental and bioinformatic approaches to investigate how naturally occurring intra-host HCV IRES diversity shapes viral translation at the level of individual mutations, complete haplotypes, and reconstructed viral populations across different biological contexts. Using patient-derived sequence analysis, mutational studies, bicistronic reporter assays, replication-deficient full-length HCV clones, and population reconstruction, we show that IRES mutations can interact epistatically and that genotype 3a populations exhibit emergent translational properties arising from interactions among coexisting variants. These findings demonstrate that translational output is not solely determined by the dominant haplotype, providing new insight into how translation contributes to viral evolution, fitness, and adaptation.

## Results

### Nineteen distinct IRES variants were identified as consensus sequences in chronically infected patients with HCV genotypes 1a, 1b, and 3a

To investigate IRES variability in patients with chronic HCV infection, we PCR-amplified the complete IRES regions from 43 samples. Each sequence was aligned and compared to the corresponding reference for each genotype. Genotype assignment had been previously determined through NS5B phylogenetic analysis(45), revealing that 26 samples belong to genotype 1a (Gt1a), 12 to genotype 1b (Gt1b), and 5 to genotype 3a (Gt3a) (Table S1). It is important to note that, despite being the most conserved region in the HCV genome, the IRESs of the reference sequences exhibit different genetic distances between each other (Gt1a versus Gt3a = 0.08117; Gt1a versus Gt1b = 0.00631; Gt1b versus Gt3a = 0.07312).

A total of 19 distinct IRES variants were identified from the consensus sequences obtained by Sanger sequencing (Table S1), with 13 of them being unique to individual samples. Among the 12 Gt1b samples, 67% (n=8) exhibited no mutations compared to the reference sequence (strain Con1), indicating that Gt1b is the least variable genotype. In contrast, only 8% (n=2) of Gt1a IRES sequences were identical to strain H77 (reference sequence for Gt1a), while all 5 Gt3a samples exhibited changes relative to the reference strain NZL1.

Most mutations mapped to IRES domains II and III (Table S1), which mediate eIF5-dependent GTP hydrolysis, 40S recruitment, and eIF3 interaction(46, 47). As expected, no mutations were detected in the conserved G-rich triplet of subdomain IIId, consistent with its essential role in 18S rRNA binding(10, 12) (Fig. S1).

Roughly half of the mutated positions were located in paired regions (nt 107, 207, 247 and 248) and often preserved Watson-Crick pairing, whereas the remaining changes occurred in single-stranded regions (nt 119, 203, 204, 205, 214, 243, 340, and 358).

These substitutions are therefore predicted to have limited effects on global secondary structure, although they could still influence binding to ribosomes, eIF3, or ITAFs.

### Relative translation activity (RTA) is not associated with HCV genotype

To assess whether genotype-specific variation influences IRES-mediated translation, we performed *in vitro* translation assays using consensus sequences derived from each viral population (Fig. 1a). Consensus IRES sequences identified by Sanger sequencing were validated by molecular cloning and resequencing, confirming both their physical presence and their predominance within each population, hereafter referred to as “master sequences”. This approach enabled a direct comparison of the translational efficiency of representative IRES variants across genotypes. In most cases, translational efficiency was comparable to that of the commonly used reference strains H77 (subtype 1a) and Con1 (subtype 1b) (Figs. 1b and 1c). Notable exceptions included two Gt1a variants, IRES 5 (mutation A243G) and IRES 11 (mutations A119C+A204C+U248C), which showed reduced activity. In addition, three Gt3a IRES master sequences were strongly impaired, approaching the inefficient control. Notably, all three shared the mutations A119C and G203A. By contrast, Gt1a IRES 3 (A204U) and Gt3a IRES 16 (U247C) displayed enhanced activity.

**Fig 1.**
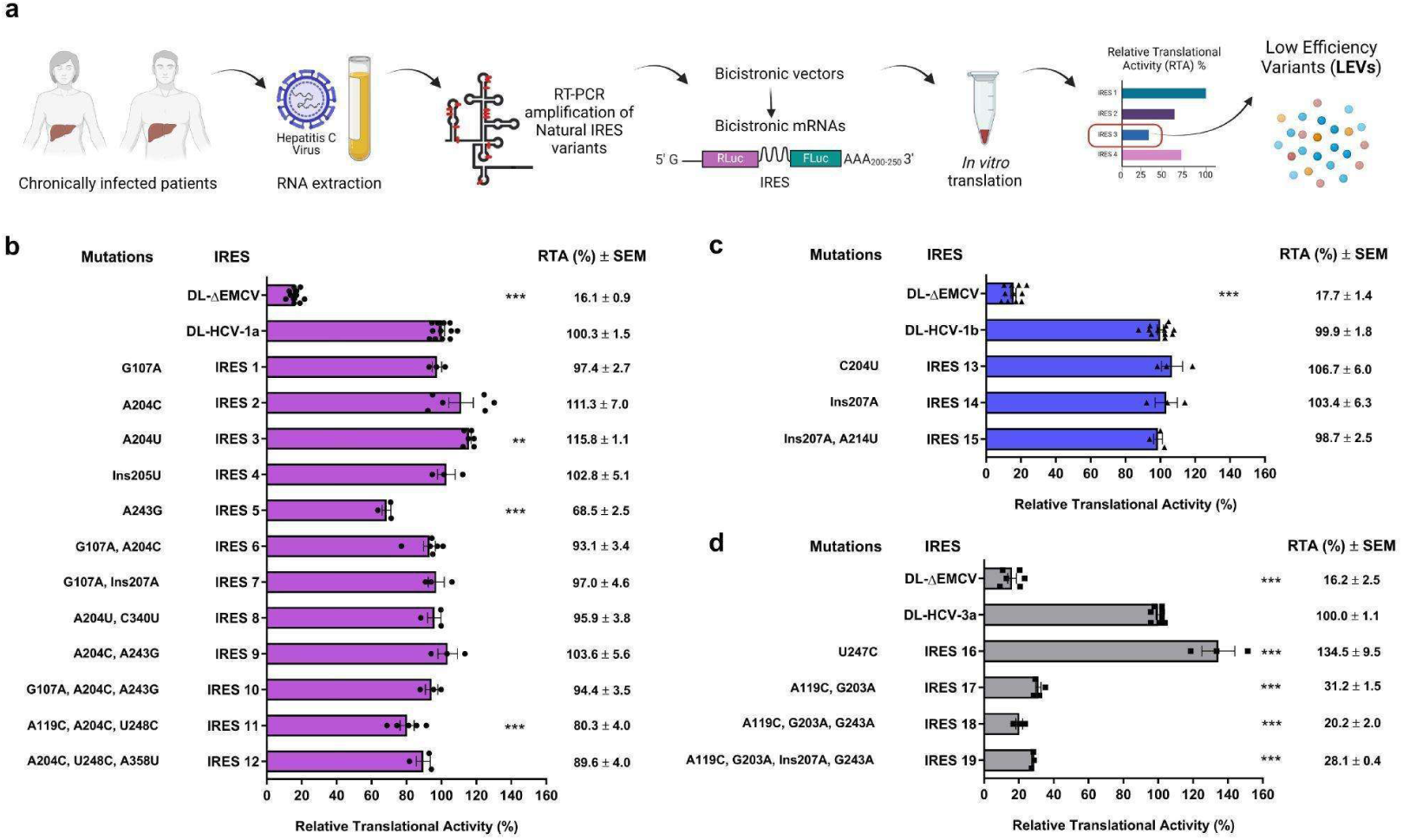
*In vitro* RTA of natural IRES variants identified as master sequences in HCV clinical isolates, analyzed using dual-luciferase constructs. IRES sequences correspond to consensus variants that were cloned into pGem-T Easy vector and subsequently subcloned into a dual-luciferase reporter vector. Natural nucleotide variations were identified through sequencing and alignment with HCV reference sequences: Gt1a (H77 strain, AF009606), Gt1b (Con1 strain, AJ238799), and Gt3a (NZL1 strain, D17763). RTA of each IRES was evaluated *in vitro* using rabbit reticulocyte lysate (RRL) by comparison with the respective positive control (DL-HCV-1a, DL-HCV-1b, and DL-HCV-3a), using DL-ΔEMCV vector as a negative control. The relative luciferase activities were measured, and the FLuc/RLuc ratio was used as the activity index, with the ratio for each positive control arbitrarily set to 100%. **(a)** Schematic representation of the *in vitro* translational assay and bicistronic reporter constructs. Generated with Biorender.com. RTAs for master sequences are shown for **(b)** Gt1a (purple bars, black circles), **(c)** Gt1b (blue bars, black triangles), and **(d)** Gt3a (grey bars, black squares).

Taken together, while specific mutations were associated with altered translation efficiency, no clear association was observed between HCV genotype and the overall RTA.

The means and standard errors (SEM) of at least three independent experiments are plotted. A one-way ANOVA test was employed to identify significant differences RTA values, followed by a Dunnett’s multiple comparison test: *** *p*<0.001, ** *p*<0.01. Significant differences were detected within each genotype group: **(b)** Gt1a, F = 89.55, df = 13; **(c)** Gt1b F = 220.5, df = 4; **(d)** Gt3a F = 259.2, df = 5.

### RTA of master IRES variants suggests epistasis at the translational level

To determine how individual mutations contributed to the phenotype of multi-mutant master sequences, we generated and tested single-point mutants using bicistronic constructs (Fig. 2). We focused on five Gt1a variants carrying multiple mutations (IRES 6, 8, 9, 10, and 11) (Fig. 2a-e) and three Gt3a variants with reduced activity (IRES 17, 18 and 19).

**Fig 2.**
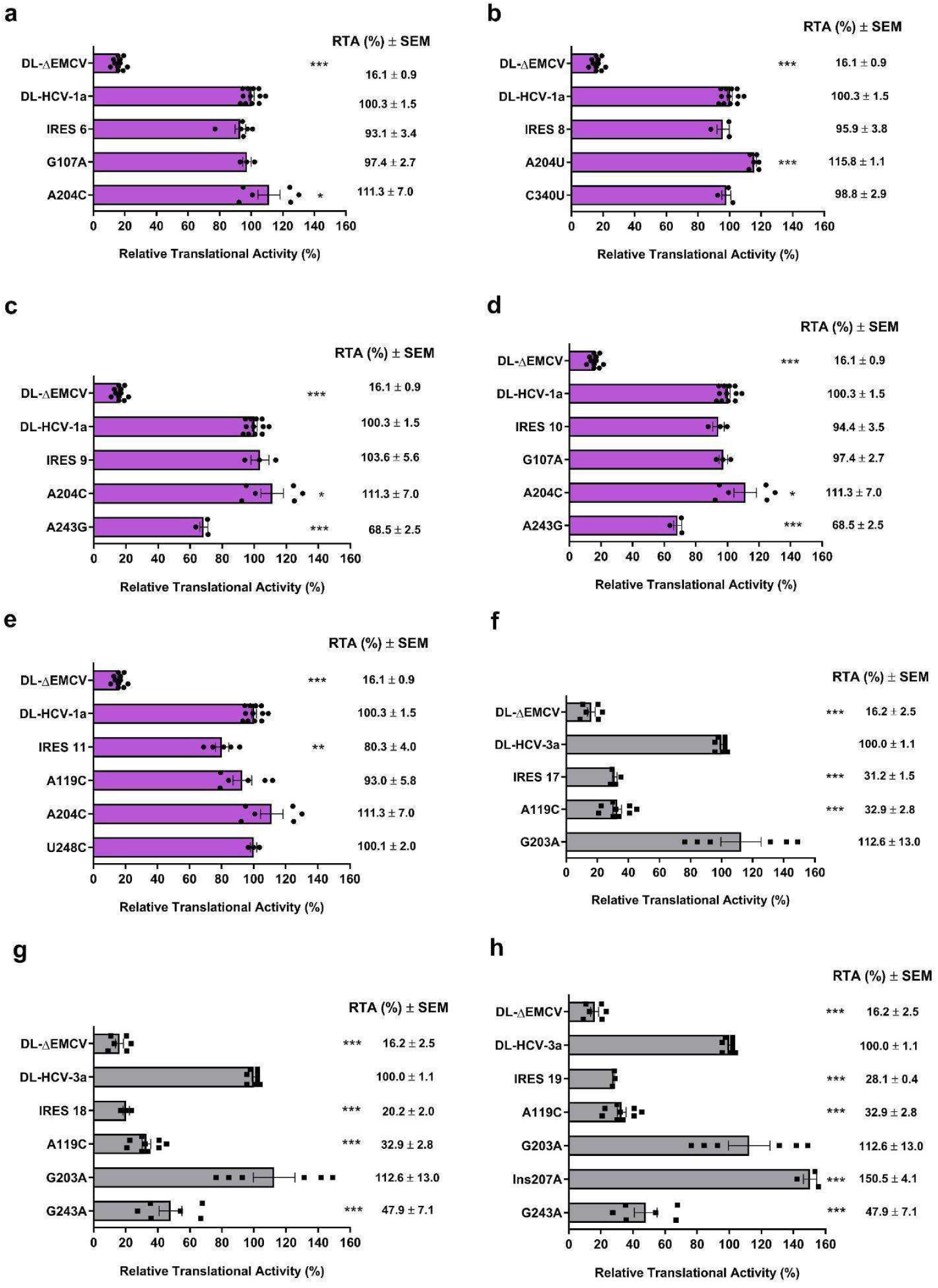
Impact of specific point mutations on the *in vitro* RTA of master IRES variants. Master IRES variants carrying multiple mutations, along with their respective single-point mutations, are presented. Controls are displayed at the top of each panel. **(a)**, **(b)**, **(c)**, **(d)**, **(e)** Gt1a IRESs: IRES 6, 8, 9, 10, and 11, respectively (purple bars, black circles). **(f)**, **(g)**, **(h)** Gt3a IRESs: IRES 17, 18, and 19, respectively (grey bars, black squares). The translational activity of each IRES was evaluated *in vitro* using RRL with DL-HCV-1a or DL-HCV-3a serving as positive controls and DL-ΔEMCV as a negative control due to its inefficient translation. Relative luciferase activities were measured, and the FLuc/RLuc ratio was used as the activity index. The ratio corresponding to each of the controls was arbitrarily designated as 100%. The plotted values represent the means of at least three independent experiments, and the bars represent SEM. Asterisks indicate IRESs with significant differences in their efficiency (one-way ANOVA followed by a Dunnett’s multiple comparison test: *** *p*<0.001, ** *p*<0.01, * *p*<0.05). **(a)** IRES 6: F = 211.2, df = 4; **(b)** IRES 8: F = 791.0, df = 4; **(c)** IRES 9: F = 203.8, df = 4; **(d)** IRES 10: F = 180.9, df = 5; **(e)** IRES 11: F = 132.7, df = 5; **(f)** IRES 17: F = 60.6, df = 4; **(g)** IRES 18: F = 46.0, df = 5; **(h)** IRES 19: F = 56.7, df = 6.

These analyses showed that the effect of a mutation depended on its sequence context. In Gt1a, changes at position 204 increased translation when tested alone but lost this effect in combination with mutations at positions 107 or 340. In IRES 9 and 10, A204C partially offset the negative effect of A243G, whereas in IRES 11 three individually neutral mutations were associated with reduced activity when combined. In Gt3a, the combination of A119C and G243A reduced translation, and the gain conferred by Ins207A alone did not rescue this defect. RNA integrity controls excluded differential degradation as a confounding factor (Fig. S2).

Our findings collectively suggest the presence of epistatic interactions at the translational level. Specifically, within Gt1a, there is evidence for antagonistic (negative) epistasis (Fig. 2a-b), since the beneficial effect of mutations in position 204 is lost when a second mutation is present. For IRES 9 and 10 (Fig. 2c-d), the translational effect of one beneficial mutation (A204C) appears to compensate for or suppress the detrimental effect of a second mutation (A243G). Furthermore, IRES 11 (Fig. 2e) may exhibit magnitude epistasis, where the collective effect of three individual neutral mutations leads to a reduction in translational efficiency. Conversely, negative synergistic epistasis may be observed in the case of Gt3a IRESs (Fig. 2g-h). Mutations A119C and G243A, both individually deleterious, are associated with an even more inefficient phenotype when combined in the same IRES sequences.

### Reference IRES sequences from different HCV genotypes exhibit substantial differences in RTA *in vitro* and *ex vivo*

We next compared the reference IRESs from genotypes 1a, 1b, and 3a using bicistronic RNAs *in vitro* (RRL) and *ex vivo* (Huh-7.5 cells) (Fig. 3). In both settings, Gt1a showed the highest activity and Gt3a the lowest, with differences becoming more pronounced in Huh-7.5 cells. Notably, translation of Gt1b and Gt3a was reduced in the cellular context. Together, these results indicate that both viral genotype and cellular environment shape HCV IRES-mediated translation.

**Fig 3.**
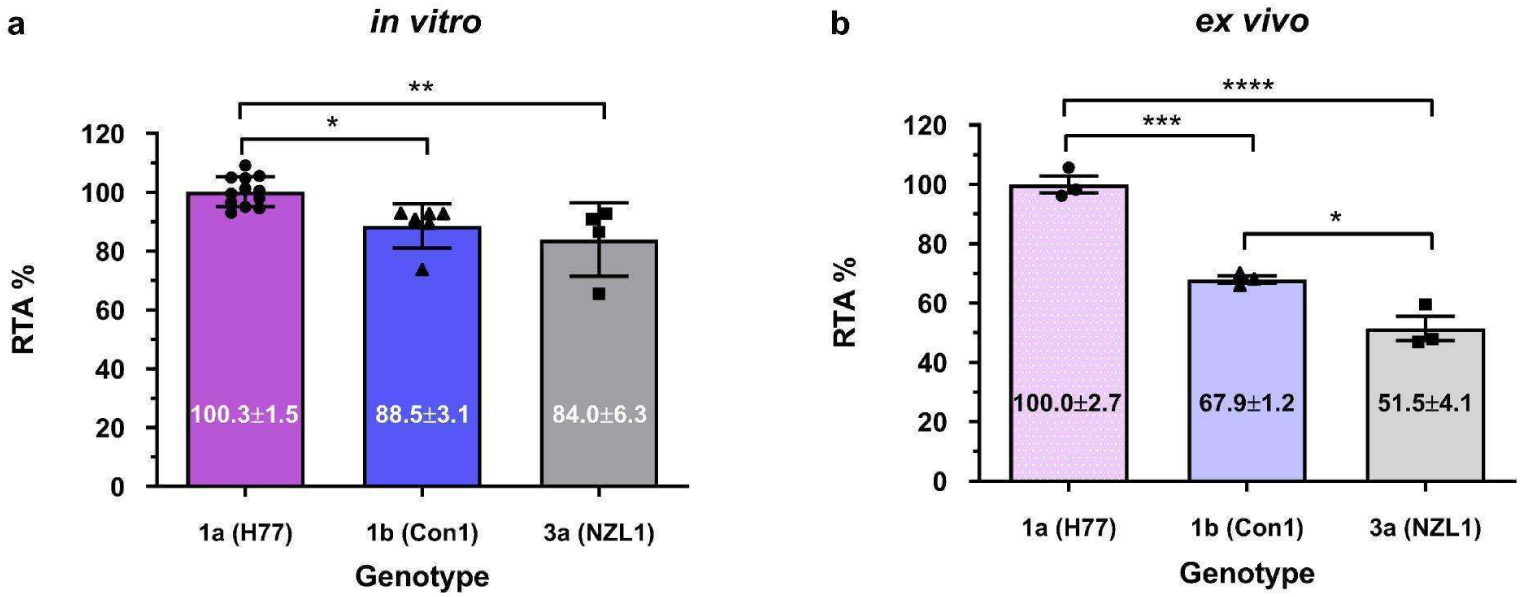
Comparative analysis of RTAs of reference IRES sequences from HCV genotypes 1a, 1b, and 3a using dual-luciferase constructs *in vitro* versus *ex vivo*. This figure compares the translational activity of reference IRES sequences from Gt1a (purple bar, black circles), Gt1b (blue bar, black triangles), and Gt3a (grey bar, black squares) in two distinct settings: *in vitro* using RRL (full colors) **(a)** and *ex vivo* in Huh-7.5 cells (faded colors) **(b)**. The relative luciferase activities were measured, and the FLuc/RLuc ratio was used as the activity index. In each case, the ratio corresponding to DL-HCV-1a was arbitrarily designated as 100%. The plotted values correspond to the means of at least three independent experiments, and the bars represent SEM. RTA and SEM values are indicated within each bar. Asterisks indicate significant differences in RTAs (one-way ANOVA, with Bonferroni correction: **** *p*<0.0001, *** *p*<0.001, ** *p*<0.01, * *p*<0.05). The test revealed significant effects among groups: (a) *in vitro* translation F = 9.629, df = 2); (b) *ex vivo* translation F = 70.05, df = 2.

### IRES genetic background and cellular context modulate RTA

Because HCV translation is influenced not only by IRES but also by downstream RNA elements, long-range RNA-RNA interactions, and cellular trans-acting factors(18, 48), we next compared bicistronic assays analyzed *in vitro* with full-length replication-deficient HCV RNAs in Huh-7.5 cells (Fig. 4a). We focused on Gt1a and Gt3a variants that showed informative phenotypes *in vitro*.

**Fig 4.**
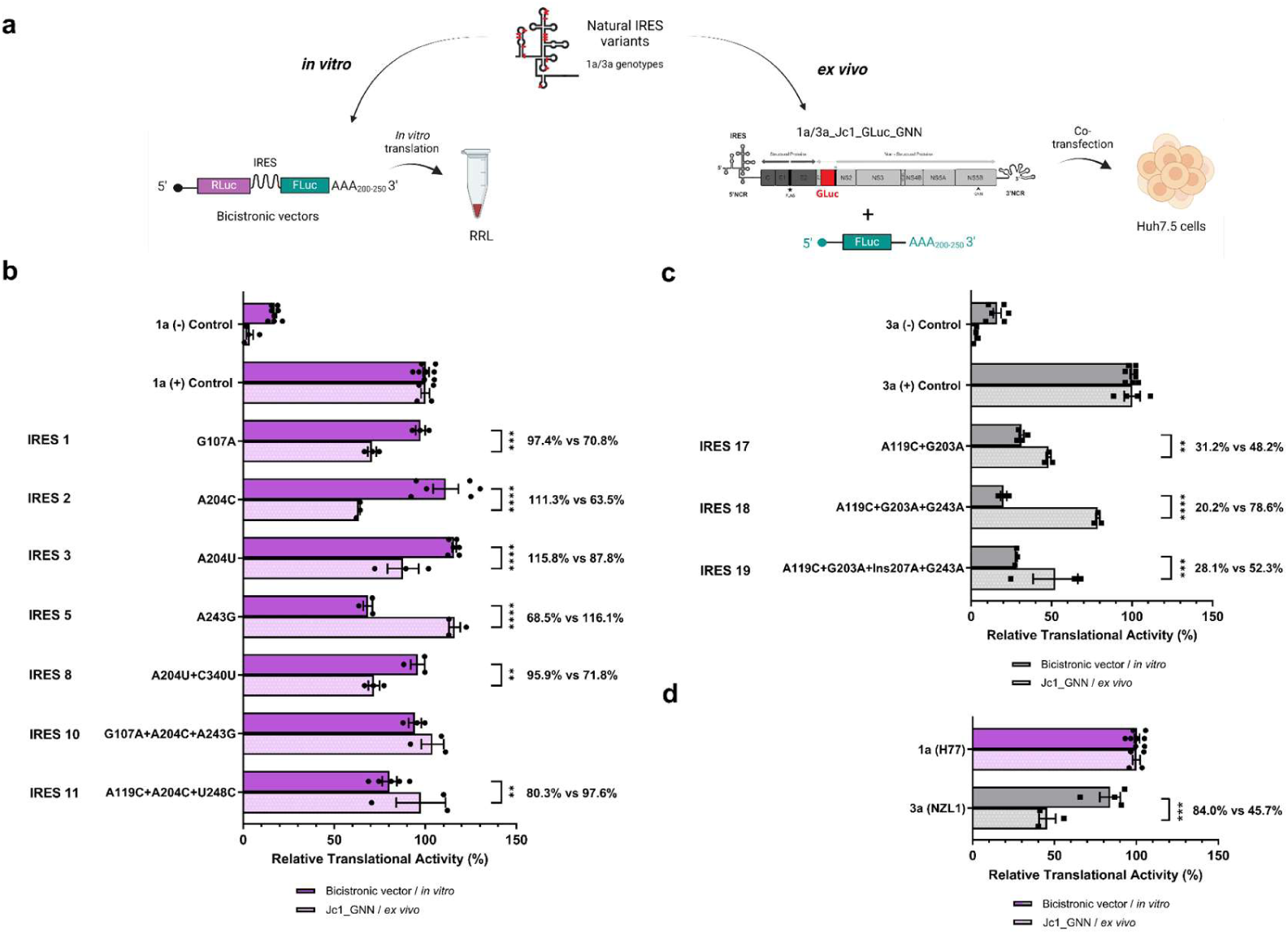
*Ex vivo* RTAs of natural IRES variants, identified as master sequences in HCV clinical isolates, analyzed in the context of replication-deficient HCV molecular clones and compared with *in vitro* assays. For *ex vivo* translational analyses, capped-mRNA encoding FLuc was co-transfected with full-length genome HCV RNA into Huh-7.5 cells. The relative translational activity of each IRES was calculated by comparing it with its respective positive control (1a-Jc1-GLuc/GNN or 3a-Jc1-GLuc/GNN), whereas mutants 1a-266+268-Jc1-GLuc/GNN and 3a-266+268-Jc1-GLuc/GNN were employed as negative controls. FLuc and GLuc activities were quantified, with the GLuc/FLuc ratio serving as the activity index. The ratio corresponding to each of the controls was arbitrarily designated as 100%. The means and SEM of at least three independent experiments are plotted. A two-way ANOVA test, with Tukey’s multiple comparisons test, was used to calculate significant differences in efficiencies between both assays: **** *p*<0.0001, *** *p*<0.001, ** *p*<0.01, * *p*<0.05. **(a)** Schematic representation of the *in vitro* bicistronic constructs and *ex vivo* full-length RNA assays. Generated with Biorender.com. **(b)** Gt1a variants (solid and faded purple bars, black circles): ANOVA interaction F = 17.62, df = 8; mutants F = 101.6, df = 8; viral/cellular context F = 13.64, df = 1; **(c)** Gt3a variants (solid and faded grey, black squares): ANOVA interaction F = 23.44, df = 4; mutants F = 189.80, df = 4; viral/cellular context F = 48.05, df = 1**; (d)** Reference genotypes: ANOVA interaction F = 27.02, df = 1; IRES reference sequence = 94.07, df = 1; viral/cellular context F = 28.15, df = 1 . In **(d)**, Gt1a was arbitrarily set as 100%.

For these *ex vivo* assays, we constructed full-length Jc1-based HCV chimeras incorporating genotype-specific 5’ non-coding regions (NCRs), and the corresponding IRES mutations. 5’NCRs sequences corresponded to the master IRES variants previously analyzed in the dual-luciferase system. These sequences were introduced into replication-deficient (GNN) Jc1 chimeras (Gt2a backbone) carrying a Gaussia luciferase (GLuc) reporter gene. The resulting chimeras enabled quantitative assessment of IRES-dependent translation in the context of full-length viral RNA.

The comparative analysis revealed that translation phenotypes were not always preserved across assay systems (Fig. 4b-c). In Gt1a, most variants behaved differently in full-length viral RNA contexts than in the bicistronic context; mutations at position 204 (A204C and A204U) generally reduced protein output *ex vivo*, whereas A243G showed the opposite effect. In Gt3a, variants carrying A119C and G203A ± G243A were more efficient *ex vivo* than predicted from bicistronic assays.

Despite these context-dependent shifts, Gt3a remained less efficient than Gt1a when the reference sequences were directly compared in the full-length system, showing an approximately 38% reduction in RTA (Fig. 4d).

To further assess context-dependent effects, we systematically evaluated point mutations *ex vivo* (Fig. S3). These analyses supported the trends observed with master sequences. In addition to the Gt1a mutants (A204C/U and A243G), A119C reduced protein output in both Gt1a and Gt3a full-length RNAs, and G203A in Gt3a displayed altered behavior *ex vivo* compared to *in vitro*.

### HCV IRES populations are characterized by multiple low-frequency variants and greater diversity in genotype 3a

To examine intra-host IRES diversity, we selected six samples whose IRES master sequences exhibited divergent *in vitro* and *ex vivo* behavior (Table S2). To this end, next-generation sequencing (NGS) and molecular cloning approaches were performed. NGS was successful in five of them (IRES 5, 8, 11, 18 and 19) with a mean coverage value ranging from 2.431X to 15.352X, with a mapping quality mean of 60 (Fig. S4 and Table S3).

We quantified within-sample diversity by Shannon entropy using both NGS reads and cloned haplotypes. The positional entropy profiles were broadly concordant between approaches (Fig. 5), supporting the robustness of the diversity estimates. Sample 05 (IRES 18) stood out as the most diverse population, consistent with prior evidence of mixed infection[49]. More generally, Gt3a samples displayed higher Shannon entropy than Gt1a, although this difference was not statistically significant (Fig. S5).

**Fig 5.**
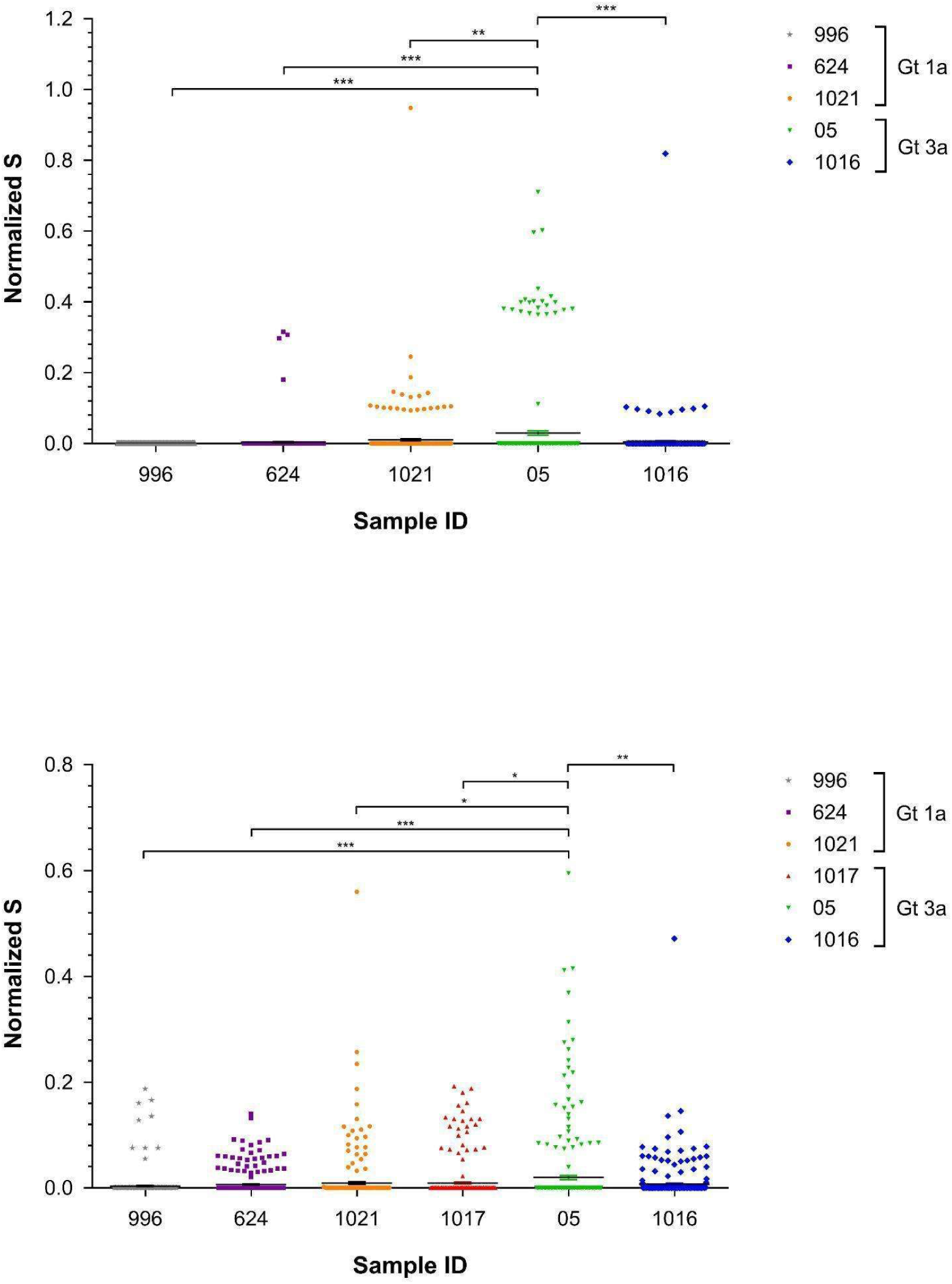
Normalized Shannon entropy per position per sample. In all cases only nucleotide variations between positions 41 and 360 of the HCV genome were considered. **(a)** Analysis conducted using NGS data. All changes exhibiting a frequency greater than 1% were included in the analysis. Shannon entropy was calculated at each position and normalized to the logarithm of the read count at that position. **(b)** Analysis conducted using molecular cloning data. All detected changes were considered. Shannon entropy was calculated with the Sequence Diversity Dynamics Analyser (DiMA https://dima.bezmialem.edu.tr./)(49) for each position and normalized to the logarithm of the clone count. The resulting scatter plots include the means and SEM of the entropy values. Significant differences in population diversity among samples were calculated using a one-way ANOVA test, with Bonferroni correction: *** *p*<0.001, ** *p*<0.01, * *p*<0.05. The test revealed significant effects among groups: **(a)** NGS data F = 11.56, df = 4); **(b)** Molecular cloning data F = 6.138, df = 5.

Cloning approaches are outlined in Table S4. Analysis of 54 to 91 clones per sample confirmed the consensus sequences as the dominant (master) haplotype and revealed multiple additional low-frequency haplotypes (Tables S4 and S5). Mixed infections detected in four out of six samples, evidenced by the presence of one to four clones corresponding to a distinct (minor) genotype, further emphasized the complexity of intra-host IRES populations.

Thus, each sample contained a dominant master sequence together with multiple minor haplotypes, with master-sequence frequencies ranging from 34% to 80% (Tables S4 and S5). The trend toward higher diversity in Gt3a was driven in part by sample 05, whose mixed infection likely explains its particularly high entropy.

### IRES population heterogeneity is a critical factor influencing translational efficiency in HCV genotype 3a

We next tested whether IRES population heterogeneity affects translational efficiency by reconstructing four intra-host populations selected for their distinct assay behavior and population complexity (Tables S2 and S4): samples 624 and 996 (Gt1a), exhibiting 20 and 11 variants, respectively, and 05 and 1017 (Gt3a), harboring 23 and 14 variants, respectively. These reconstructed populations included the dominant master haplotype plus low-frequency variants present in 1-6% of the population.

Variants incompatible with the Jc1 backbone (due to mutations within the core region differing between Gt1a/3a and 2a), those that could not be cloned, or those belonging to a minor genotype in mixed infections were excluded, and frequencies were renormalized accordingly (Table S5).

Specifically, three variants from sample 05, and one variant each from samples 1017 and 996 were excluded on these grounds. In addition, one variant from sample 624, containing a cytosine insertion at position 127, that generated an extended homopolymeric stretch (5′ CCCCCCCCCUCCC 3′), could not be successfully cloned despite repeated attempts and was therefore omitted. Exclusion of minor-genotype variants in mixed infections affected samples 624 and 1017 (one variant each) and sample 05 (four variants). The total number of variants for each population and their corresponding adjusted frequencies are shown in Table S5.

Reconstituting the viral population had little effect on translation for the two Gt1a samples (Fig. 6a-b). In contrast, reconstructed Gt3a populations translated significantly better than their corresponding master sequence alone (Fig. 6c-d). These results indicate that intra-host IRES heterogeneity can modulate translation at the population level in Gt3a, whereas this effect was not detected for Gt1a under the conditions tested.

**Fig 6.**
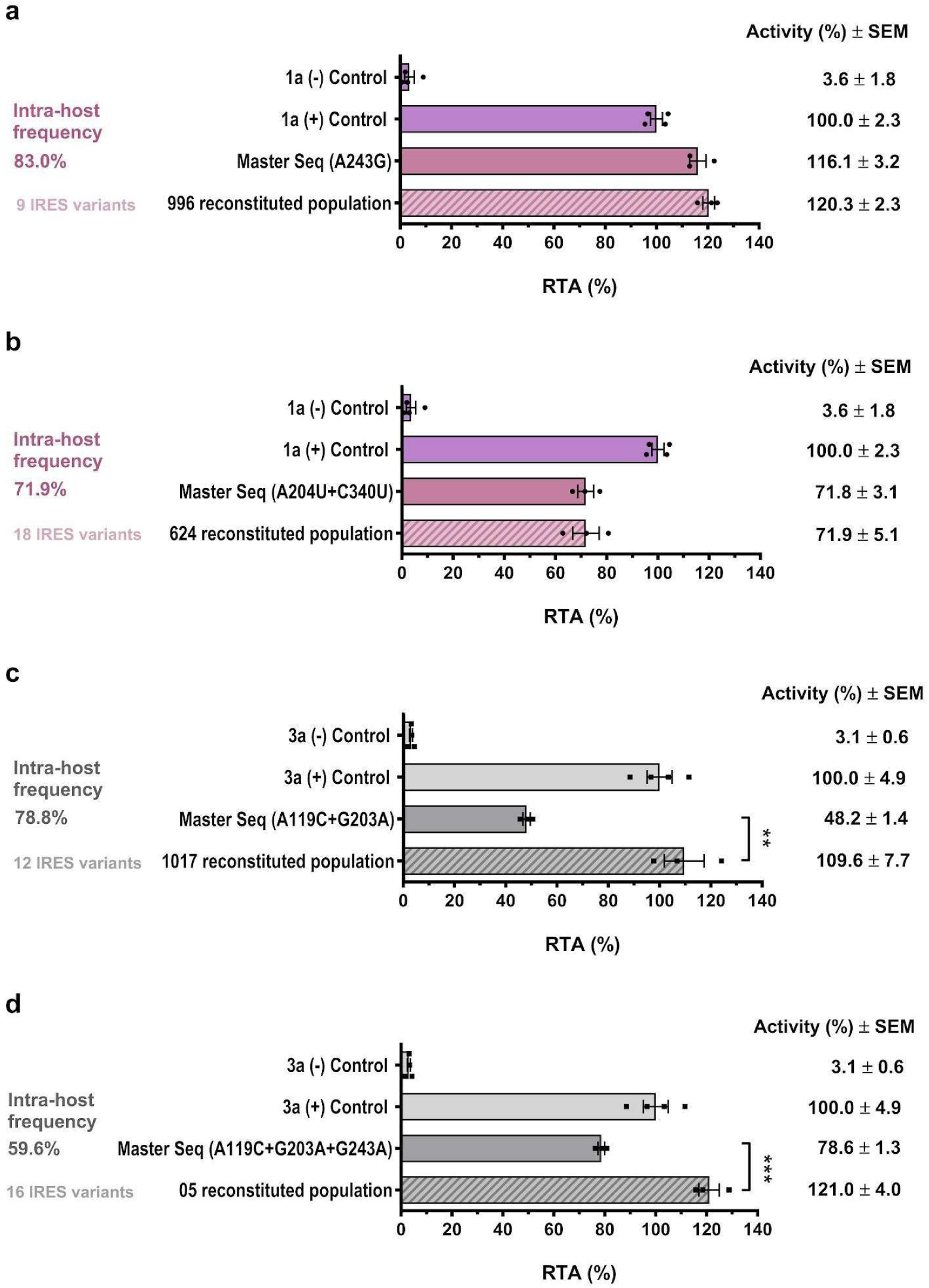
*Ex vivo* RTAs of reconstituted IRES populations versus their corresponding master sequences. *Ex vivo* experiments were conducted with replication-deficient (GNN) Jc1-GLuc (Gt2a) chimeras harboring 5’NCR regions of Gt1a or Gt3a. HCV RNAs were co-transfected with a capped mRNA encoding FLuc into Huh-7.5 cells. Master sequence IRESs (dark colored bars) were transfected as single sequences and compared to each corresponding reconstituted population (colored hatched bars), where the adjusted frequency of each variant was considered (S4 Table). The relative translational activity was evaluated by comparing each construct with its respective positive control (1a-Jc1-GLuc/GNN or 3a-Jc1-GLuc/GNN), and mutants 1a-266+268-Jc1-GLuc/GNN or 3a-266+268-Jc1-GLuc/GNN were employed as negative controls (inefficient). Positive and negative controls are shown in faded purple (Gt1a) and faded grey bars (Gt3a). FLuc and GLuc activities were quantified, with the GLuc/FLuc ratio serving as the activity index. The ratio corresponding to each of the positive controls was arbitrarily designated as 100%. The means and SEM of at least three independent experiments are plotted. A two-tailed unpaired t-test was used to calculate significant differences in RTA between master sequences and reconstituted populations. ** *p*<0.01, *** *p*<0.001. The percentages of the master IRES sequence within each population are indicated in orange, whereas the total number of variants (haplotypes) of each population are indicated in blue. The different panels illustrate the results of IRESs from different samples: **(a)** 996 (Gt1a) – t-value = 1.074, degrees of freedom = 4, **(b)** 624 (Gt1a) – t-value = 0.00778, degrees of freedom = 4, **(c)** 1017 (Gt3a) – t-value = 7.807, degrees of freedom = 4, and **(d)** 05 (Gt3a) – t-value = 10.13, degrees of freedom = 4.

## Discussion

Our study addresses how naturally occurring HCV IRES variation affects translation at both the haplotype and population levels, revealing that translation output is determined not only by intrinsic activity of individual IRES variants but also by epistatic interactions, assay context, and, in Gt3a, the composition of the broader intra-host viral population.

Previous work on virus intra-host variability has focused mainly on coding regions(50–53), whereas the consequences of IRES variation have remained unclear. Earlier studies relating IRES sequence to interferon-based therapy outcomes and HCV infection parameters reached inconsistent conclusions(54–58), and the contribution of intra-host IRES heterogeneity to translation has only been minimally explored(43) and exclusively by means of bicistronic vectors.

Here, we address this gap by combining patient-derived sequences, bicistronic assays, full-length replication-deficient HCV clones, NGS, cloning, and population reconstruction. This integrative approach enabled us to directly compare individual haplotypes with the populations from which they originated, and extends previous studies that relied exclusively on bicistronic reporter systems by evaluating naturally occurring IRES diversity in a more biologically relevant genomic context.

Several key conclusions emerge from our results. First, low-RTA master variants were identified in more than one genotype, arguing against a simple genotype-level explanation for reduced translation. Nevertheless, clear genotype-specific trends were observed: Gt1b appeared comparatively conserved, whereas Gt3a displayed the greatest sequence diversity and included the most translationally impaired master variants (Figs. 1 and 5).

Second, our mutational analyses demonstrate that IRES activity is shaped by intragenic epistasis rather than by isolated mutations (Fig. 2), indicating that the functional consequences of naturally occurring IRES variation cannot be predicted from individual substitutions in isolation. In Gt1a, some apparently beneficial changes lost their effect when combined with a second mutation, whereas in other cases one mutation buffered the effect of another. In Gt3a, combinations of individually deleterious mutations yielded even lower activity.

*Ex vivo* analyses further revealed context-dependent genetic suppression, in which combinations that appeared deleterious in bicistronic assays partially or fully recovered activity in the full-length viral RNA context (Fig. S3).

Importantly, these epistatic effects are unlikely to arise from major alterations in the predicted RNA secondary structure. Modeling of domain III revealed only modest differences in free energy and predicted interactions (Fig. S6), although key Gt3a changes could not be fully evaluated without experimentally constrained folding data, such as SHAPE(59–62). The underlying mechanisms may therefore involve local conformational effects or altered RNA-protein and RNA-RNA interactions, rather than extensive structural rearrangements.

The comparison between bicistronic and full-length systems is one of the main strengths of this study. Several IRES variants exhibited discordant behavior across assays, and genotype-specific differences were more pronounced in Huh-7.5 cells and when including the full-genome context, indicating that both cellular environment and genomic architecture shape IRES function (Figs. 3 and 4, respectively). These observations highlight an important limitation of bicistronic *in vitro* assays: although widely used, they do not fully recapitulate the complexity of translation within the native viral RNA, which had not been addressed so far. Previous studies demonstrated that IRES activity may also vary among cell types, but those analyses were likewise restricted to bicistronic constructs(38, 41, 42).

A central finding of our work is that intra-host IRES heterogeneity influences translational efficiency at the population level particularly in HCV Gt3a (Fig. 6). Reconstituted Gt3a viral populations consistently exhibited higher translational efficiency than their corresponding master haplotypes, in some cases approaching the activity of reference sequences. In contrast, no comparable effect was observed for Gt1a under the conditions tested. These findings indicated that, in Gt3a, translational output is not solely determined by the dominant sequence but instead emerges from the combined behavior of coexisting variants.

These observations substantially extend previous studies describing the collective effect of IRES diversity, which were restricted to bicistronic reporter assays(43). By analyzing naturally occurring IRES variants in replication-deficient full-length HCV genomes in Huh-7.5 cells, our approach reproduces more closely natural infection conditions(15, 17, 30, 63–65). Because these constructs do not produce progeny, they avoid confounding effects of replication and selection, which may arise from the use of replicons-based systems(35), providing a more direct measure of translation itself. Our findings therefore, support a model in which viral populations, rather than individual master sequences, constitute the relevant functional unit governing translational output.

Several non-mutually exclusive mechanisms may explain this population-level effect (Fig. 7). One potential mechanism is trans-complementation, whereby poorly-translating variants benefit from more efficient genomes within the same cell; a phenomenon previously described in members of the *Picornaviridae* family(66–68). Alternatively, heterogeneous IRES RNA structures may cooperate by promoting more efficient recruitment of translation factors, stabilization of translation-competent ribonucleoprotein complexes, or transfer of a functional initiation complex from a functional IRES element to a defective IRES structure(69). Although these mechanisms remain to be experimentally demonstrated in HCV, they provide biologically plausible explanations for the cooperative translational behavior observed in Gt3a populations.

**Figure 7.**
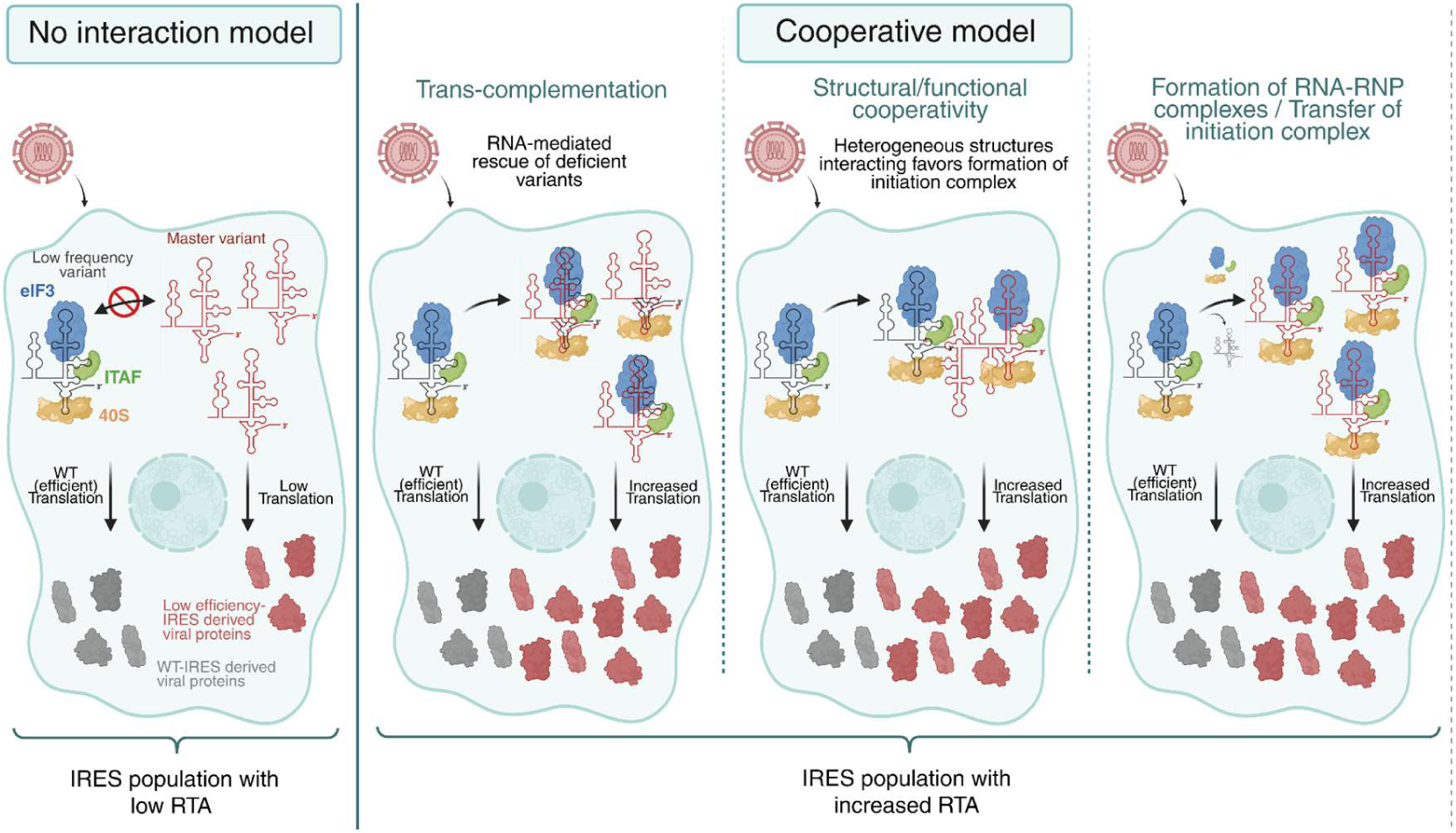
Alternative models of molecular mechanisms resulting in cooperative interactions in HCV genotype 3a. Schematic representation of possible interaction models within an IRES variant population that could explain the cooperative increase in RTA observed in HCV Gt3a. In the “No interaction” model (left, and model 1, Fig. S7a), a dominant master variant exhibits low translational efficiency, and rare wild-type (WT)-like variants fail to significantly influence overall RTA, resulting in reduced viral protein output. In contrast, a “Cooperative model” may be attributable to a variety of mechanisms (e.g., models 2 and 3, Fig. S7b-c). The “Trans-complementation” mechanism proposes that more efficient variants within the same cell can rescue poorly translating variants through RNA-mediated mechanisms, thereby enhancing total translation (middle left). The structural/functional cooperativity mechanism suggests that heterogeneous IRES structures interact to promote the assembly of translation initiation complexes (middle right). A third potential mechanism involves the formation of RNA–RNP complexes or the transfer of initiation complexes, in which stable ribonucleoprotein assemblies or functional initiation complexes formed on efficient IRES elements promote translation from otherwise defective variants (right). Relative protein translation of these alternative scenarios, with respect to model 1, was modeled using competition coefficients that account for IRES-IRES (α) and/or protein–cofactor (λ) interactions. At equilibrium, the total protein output with respect to model 1 (*Δ*PT) depends on the interacting coefficients. Negative competition coefficients (α < 0, λ < 0) suggest cooperative translation (Fig. S8-S9).

To explore whether cooperative interactions among IRES variants could, in principle, account for our experimental observations, we developed a simple resource-consumer model in which multiple IRES variants compete for limiting translation cofactors (Fig. 7, Fig. S7). In this framework, interactions among variants can shift the system from purely competitive behavior toward net cooperation, analogous to ecological systems in which consumer interactions become mutualistic(70) (Fig. 7, Fig. S7).

Although simplified, the model supports the idea that cooperative interactions among IRES variants are sufficient in principle to increase translation, and helps reconcile previous discordant reports on the functional significance of IRES variability(13, 35, 40–42, 66–71).

One interpretation of these findings is that IRES heterogeneity may act as a bet-hedging strategy, allowing the viral population to maintain translation across variable intracellular environments even when the dominant haplotype is not itself optimal. Our data also suggest that Gt3a populations may be more translationally robust than Gt1a, because increased diversity was associated with recovery of population-level output despite low-efficiency master sequences(72). Whether this enhanced robustness contributes to biological properties historically associated with Gt3a, including differential treatment response(73–75) remains speculative and will require direct testing in clinically annotated cohorts.

This study has several limitations. The number of reconstructed viral populations was modest, and some naturally occurring variants could not be included for technical reasons. The relatively small number of Gt3a samples limited statistical power and precluded subgroup analyses, including comparisons between responders and relapsers. In addition, because all patients were chronically infected, we could not address the effects of infection stage, and incomplete clinical metadata prevented a formal analysis of the relationship between translational activity and treatment outcome, despite previous reports suggesting such an association(57). Future studies including larger longitudinal cohorts and additional cellular models, will be important to determine the broader biological and clinical relevance of these observations.

In summary, our results demonstrate that the translational phenotype of naturally occurring HCV IRES variants is an emergent property shaped by epistatic interactions, genomic context, and the composition of intra-host viral populations. Bicistronic reporter assays and replication-deficient full-length systems capture overlapping but non-identical aspects of IRES functions, emphasizing the importance of studying regulatory RNA elements within their native genomic context. Furthermore, the enhanced translational output observed in reconstructed Gt3a populations, together with our quantitative modeling, supports the concept that cooperative interactions among coexisting viral variants can modulate translation at the population level.

Altogether, our findings expand the current view of HCV IRES function by showing that naturally occurring regulatory RNA variation cannot be fully understood by analyzing individual mutations or dominant haplotypes in isolation. Instead, the functional consequences of IRES diversity emerge from interactions occurring across multiple biological levels, including mutation combinations, genomic context, and the genetic composition of viral populations. More broadly, these results suggest that population-level properties of non-coding regulatory elements may represent an underappreciated determinant of RNA virus phenotypes and evolution, with implications extending beyond HCV to other genetically diverse RNA viruses.

## Materials and methods

### Clinical samples and ethical considerations

We performed an observational and cross-sectional study in which 43 serum samples were collected from chronically infected HCV patients. All patients were 18 years or older and serologically negative for Hepatitis B virus. Samples were collected between December 18th, 2014, and December 20th, 2016, in the Hospital de Clínicas, a public university hospital in Montevideo, Uruguay. This study was conducted according to national and international ethical guidelines (good clinical practice and the Declaration of Helsinki) and was approved by the institutional ethics committee. Written informed consent was obtained from all participants, and personal information was kept strictly confidential, with access limited to medical personnel. Additionally, demographic data and virological parameters, including viral load, antiviral therapy, and response, were recorded (Table S6).

### Genotype determination of clinical samples

HCV genotyping of clinical samples was previously conducted and is detailed in Echeverría et al. (2024)(45). Briefly, RNA was extracted from serum samples using the QIAamp Viral RNA Mini Kit (QIAGEN, Hilden, Germany), followed by cDNA synthesis using Superscript II reverse transcriptase (Invitrogen Life Technologies, Carlsbad, CA, USA) according to the manufacturer’s instructions. A 386 bp conserved region of the NS5B polymerase gene was then amplified using a hemi-nested PCR protocol, and the resulting sequences were phylogenetically analyzed. The genotyping results are shown in Table S6.

### One-step PCR of the IRES region and sequence analyses

Extracted RNA was subjected to a one-step PCR reaction using previously described primers that amplify the entire IRES region (370 bp encompassing nucleotides 14 to 383 of the genome, according to H77 strain numbering - NCBI accession number AF009606)(13). The PCR mix and reaction conditions have been reported in detail(45). Since both primers contain recognition sites for the restriction enzymes *Xho*I and *Eco*RI to facilitate cloning and subcloning the products, the generated PCR amplicon is 391 bp long. IRES products were visualized on 2% agarose gels stained with SYBR Safe DNA gel stain (Invitrogen, Madison, USA), purified, and sent to Macrogen Inc. (Korea) for bidirectional sequencing. Sequences were edited using Seqman software implemented in DNAStar 5.01 (DNASTAR, Madison, USA) to generate a consensus sequence for each IRES, representing the most frequent nucleotide at each position across the set of sequences derived from each host. Mutation detection was performed by comparison against 3 reference sequences corresponding to Gt1a (H77 strain, NCBI accession number: AF009606), Gt1b (Con1 strain, NCBI accession number: AJ238799), and Gt3a (NZL1 strain, NCBI accession number: D17763), according to the genotype of the sample as determined by the NS5B phylogeny.

### Isolation of natural IRES variants and cloning in bicistronic vectors

In order to analyze the translational efficiency of different IRES variants (haplotypes), a selection of IRESs with mutations in their consensus sequences with respect to the reference sequences was made, and their master IRES variants were isolated (the most abundant variant or the dominant sequence within a viral population). Additionally, for selected samples, one or two IRES haplotypes that appeared as low-frequency variants (present in 1-6% in each viral population) were also isolated for further analysis. To isolate these variants, IRES PCR products were adenylated with 5U Taq DNA Polymerase, recombinant (Fermentas, Vilnius, Lithuania), and 0.5µM dATP and subsequently cloned into pGem-T-Easy vector (Promega, Madison, USA). A molar ratio of 3:1 (insert:vector) was used according to the manufacturer’s instructions. Next, 10µL of the ligation mixture was transformed into chemo-competent DH5α *E. coli* cells (heat shock 42°C - 30sec). The bacteria were then plated on ampicillin-IPTG-X-gal containing plates and grown overnight at 37°C. After incubation, 15-20 white colonies were used to make 2mL starter cultures and grown overnight at 37°C - 220 rpm.

Plasmids were then extracted using the PureLink Quick Plasmid MiniPrep (Invitrogen, Vilnius, Lithuania). Insert verification was performed by PCR amplification using universal primers M13F and M13R. Positive clones were sequence-verified by Sanger sequencing (Macrogen, Korea).

A clone representative of the master sequence (matching the consensus sequence) was subcloned into bicistronic vectors (DL-HCV) encoding Renilla luciferase (RLuc) as the first cistron and Firefly luciferase (FLuc) as the second cistron, separated by an intergenic region into which each IRES variant was cloned (flanked by *Xho*I and *Eco*RI restriction sites). We used the DL-HCV-1b vector kindly provided by Dr. Marcelo López-Lastra (Pontificia Universidad Católica de Chile, Chile)(13) as a positive control for Gt1b sequences, and constructed positive controls for Gt1a and Gt3a by either subcloning an IRES sequence without any mutation (compared to the Gt1a reference strain) or by site-directed mutagenesis reversing a mutation present in a Gt3a clone. The plasmid DL-ΔEMCV, used as an inefficient translational control, was also provided by Dr. Marcelo López-Lastra and corresponds to the RLuc-delta EMCV-FLuc vector, which lacks an active IRES element(76). A schematic representation of the assays and the bicistronic vectors used in this study is depicted in Fig. 1a.

### Generation of IRES single mutants in bicistronic vectors

Since some Gt1a and Gt3a IRESs with multiple mutations showed low translational activity *in vitro*, they were selected to dissect the effect of each mutation on translation. Therefore, single mutants were generated by site-directed mutagenesis using overlapping primers with the desired mutations at the center of each primer. The primers used are listed in Table S7 and were designed using Primer X Software (https://www.bioinformatics.org/primerx/index.htm). Briefly, PCR was performed using 1U Phusion High Fidelity polymerase (Thermo Scientific, USA), 1X buffer (including MgCl_2_), 0.25mM dNTPs, 0.80µM of each mutagenesis primer, 500ng of bicistronic reference vector, and DEPC water to a final volume of 40µL. The PCR cycle consisted of a pre-denaturation step at 98°C – 5min, 35 cycles of 98°C – 30sec, 60°C – 30sec, 68°C – 4.5min, followed by a final extension at 72°C – 10min. Parental DNA was then degraded by digestion with 20U of *DpnI* (New England Biolabs, Ipswich, MA, USA), according to the manufacturer’s instructions. After verifying correct amplification and parental DNA digestion by electrophoresis, 10µL of the unpurified digested product was used to transform chemo-competent DH5α *E. coli* bacteria according to the protocol described above. After plasmid extraction, all constructs were verified by Sanger sequencing (Macrogen, Korea).

### *In vitro* transcription of bicistronic RNAs

Bicistronic plasmids were linearized with *BamHI* (Invitrogen, Life Technologies, Vilnius, Lithuania), precipitated with 1/10 volume of sodium acetate (3M, pH 5.2) and 2.5 volumes of cold ethanol, and then used for *in vitro* transcription. Capped and polyA-tailed mRNAs were synthesized and purified using the HiScribe T7 ARCA mRNA Kit with tailing (New England Biolabs, Ipswich, MA, USA) and LiCl, respectively, according to the manufacturer’s specifications. RNA concentrations were measured spectrophotometrically, and RNA integrity was verified by electrophoresis on non-denaturing agarose gels. Before loading the gels, 1μg of RNA was mixed with 1X RNA Loading Dye (New England Biolabs, Ipswich, MA, USA), pre-heated at 70°C for 10 min, and then cooled on ice to remove secondary structures.

### *In vitro* translation

*In vitro* translation was performed in nuclease-treated rabbit reticulocyte lysate (Promega, Madison, USA), with optimized salt conditions(77). Reactions were performed in a final volume of 25μL, including 25ng of purified *in vitro* transcribed RNA. Incubation was carried out at 30°C for 90 min. RLuc and FLuc activities were measured using the Dual-Luciferase Reporter Assay System (Promega, Madison, USA) on a plate LUMIstar OPTIMA luminometer (BMG Labtech) according to the manufacturer’s protocol. The FLuc/RLuc ratio was used as an index of IRES activity, and the mean of this index for each reference sequence (DL-HCV-1a, 1b, or 3a) was considered as 100%. In all cases, at least 3 independent experiments with 2 technical replicates each were performed.

### Analysis of intra-host IRES population diversity

To analyze the intra-host diversity of the IRES population, we selected 6 samples that had IRES sequences with inefficient translation activities (low efficiency variants present either as master or low frequency). Three of the selected samples belonged to Gt1a (624, 996, 1021) and three to Gt3a (05, 1016, 1017). To this end, two distinct approaches were employed: cloning and next-generation sequencing.

For the cloning approach, the pCR™2.1-TOPO™-TA vector (Invitrogen, Carlsbad, CA, USA) and chemically competent Top10 *E. coli* bacteria were utilized, as previously described(45). In summary, the IRES PCR products were adenylated and purified prior to the transformation protocol. The transformation protocol was executed following a heat shock step of 30 seconds at 42°C. Subsequently, 250 µL of SOC medium was added, and the tubes were incubated for 1 hour with shaking to facilitate bacterial recovery. Next, 50 µL of each transformation was added to pre-heated LB plates containing ampicillin and X-gal, and the plates were incubated overnight at 37°C. Thereafter, 50–150 white colonies were picked for colony PCR insert verification and subculturing into 96-well plates containing LB agar plus ampicillin. These were then sent to Eurofins Genomics (Ebersberg, Germany) for PlateSeq sequencing. Sequences were then processed using Seqman software implemented in DNAStar 5.01 (DNASTAR, Madison, USA) and compared to 1a or 3a reference sequences (strain H77 - accession number AF009606 and strain NZL1 - accession number D17763, respectively) to analyze intra-host genetic variability.

For next-generation sequencing, libraries were prepared using the TruSeq Nano DNA LT Library Prep Kit (Illumina, San Diego, CA, USA), according to the manufacturer’s instructions. The quantification of IRES PCR products was conducted using the Qubit 2.0 Fluorometer dsDNA HS Assay (Life Technologies, Eugene, OR, USA) prior to their utilization. Then, a paired-end (2×250) sequencing run was performed with MiSeq Reagent Kit v2 (500 cycle) (Illumina, San Diego, CA, USA) on a MiSeq instrument (Illumina) from the sequencing service of the Institut Pasteur Montevideo. Data pre-processing, alignment, and visualization were done in the Galaxy web platform (http://usegalaxy.org)(78). The raw data underwent analysis using the FastQC program(79). To address potential contaminants, the Trimmomatic tool was employed to remove them(80). This entailed the implementation of a sliding window (4nt) algorithm, which was utilized to remove adapters, primer sequences, and all reads exhibiting an average quality score below 25 (Q<25). Subsequently, reads with a length less than 36 nucleotides were removed by the MINLEN algorithm. A quality check was performed using the FastQC program. The IRES consensus sequences, previously obtained by Sanger methodology, were employed as reference sequences for assembly. The consensus sequences were then mapped using the BWA-MEM software(81). The coverage profiles of each sample were evaluated using the Qualimap 2 tool(82). Variant calling was performed using the MPileUp algorithm from SAMTools(83). Analysis was conducted only on nucleotide positions with variant frequencies above 1%. To measure the diversity or uncertainty within the intra-host population, the Shannon entropy was calculated for each IRES position from nucleotide 41 to 360 of the HCV genome (according to H77 strain numbering). Shannon entropy measures genetic diversity by quantifying the degree of variation or randomness in the distribution of different viral genomes within a population(84). For NGS data, the entropy was calculated using custom scripts developed in Python for each position and normalized by dividing by the logarithm of the read count at that position. For molecular cloning data, it was calculated from multifasta alignments of clones from each sample using the Sequence Diversity Dynamics Analyser (DiMA, https://dima.bezmialem.edu.tr./)(49). Entropy was calculated for each position (k-mer = 1) and normalized by the logarithm of the number of clones for each sample. This normalization process enabled the entropy to be scaled by the sequencing or cloning depth, thereby controlling for potential sampling bias.

### Huh-7.5 cell line and culture conditions

Huh-7.5 cell line(85), a highly permissive line derived from parental Huh-7 hepatoma cells, was kindly provided by Charles M. Rice from The Rockefeller University (New York, USA). Cells were cultured in Dulbecco’s modified Eagle’s medium (DMEM) (Gibco, Paisley, UK), supplemented with 5% heat-inactivated fetal bovine serum (FBS) (Hyclone, Thermo Scientific, Utah, USA), 100U/mL penicillin and 100 μg/mL streptomycin (Gibco/Invitrogen), and 0.1mM non-essential amino acids (NEAA) (Invitrogen, Life Technologies, Carlsbad, CA), with 5% CO2 at 37 °C.

### Full-length HCV replicons

Plasmid pJc1FLAG(p7-nsGLuc2A)/GNN is a GLuc reporter replication-deficient GNN replicon (GDD catalytic motif of NS5B mutated to GNN), harboring an intergenotypic J6/JFH HCV Gt2a strain(86). This plasmid was used as a backbone to generate reference chimera plasmids containing 5’NCR regions of Gt1a (H77 strain) or Gt3a (NZL1 strain) by overlap-extension PCR using Phusion High Fidelity polymerase (Thermo Scientific, USA). A schematic representation of the full-length HCV replicons used in this study is depicted in Fig. 4a. The 5’NCR regions were obtained by PCR from sub genomic replicons H77/SG-Feo (L+8) - Gt1a(87) and S52/SG-Feo - Gt3a(88).

### Full-length HCV replicons with mutated IRES

Subsequent to the generation of the reference plasmids, translationally inefficient IRES were constructed by introducing mutations via overlap-extension PCR, employing the same primers listed in Table S7. Each mutation was introduced using the reverse primer for PCR1 and the forward primer for PCR2. Additionally, primers M13F (5’ GTAAAACGACGGCCAGT 3’, which anneals upstream of the T7 promoter and the *Eco*RI restriction site) and PCR2_R (5’ CATCGCGTACGCCAAGATCATG 3’, which hybridizes to E1 at the *Bsi*WI restriction site) were employed as PCR1 forward and PCR2 reverse primers, respectively. For those IRES sequences that contained multiple mutations, they were sequentially introduced.

### IRES population reconstruction

Four IRES populations, comprising a varying number of haplotypes and exhibiting differential abundance of the master sequence, were selected for further translational studies (two of Gt1a — samples 624 and 996, and two of Gt3a — samples 005 and 1017). To facilitate the reconstruction of each of these intra-host populations, a mutagenesis approach was employed using the In Phusion HD EcoDry Cloning Kit (Takara Bio, USA), per the manufacturer’s instructions. The result of this process was the creation of 55 IRES haplotypes. To that end, a linear plasmid was generated by PCR amplification using Q5® High-Fidelity DNA Polymerase (New England Biolabs) and primers BB_Jc1_F (5’ CGCCCACAAGACGTTAAGTTTCC 3’) and BB_Jc1_R1 (5’ CACTGGCCGTCGTTTTACAACG 3’), encompassing the core region of the HCV genome up to the T7 promoter region. The inserts (IRESs, variants, or haplotypes) were designed as gBlocks Gene Fragments of approximately 450 base pairs (Integrated DNA Technologies) covering the full 5’NCR region, the first 71 nucleotides from the core, and 43 nucleotides from the plasmid backbone. The insert ends feature a 15-base pair overlap that is complementary to the vector ends. The sequences of the gBlocks are available upon request. Before *in vitro* transcription, the sequence of each plasmid was verified by Sanger sequencing (Macrogen Korea).

### *In vitro* transcription of HCV full-genome constructs

HCV chimeric RNAs were *in vitro* transcribed from *Xba*I linearized DNA plasmids using the T7 RiboMAX™ Large Scale RNA Production System (Promega). Then, the RNA was treated with RQ1 DNase (Promega) at 37°C for 15 min and purified following the reaction clean-up protocol from the Quick RNA Miniprep Plus Kit (Zymo Research). The concentrations and integrity of the RNA were determined as previously described for bicistronic RNAs.

### Bicistronic RNA transfection in Huh-7.5 cells for *ex vivo* translational studies

For transfection of bicistronic RNAs for *ex vivo* translational studies, 100ng of each capped and polyadenylated bicistronic RNA were mixed with 0.5µl Lipofectamine 2000 (Life Technologies) in 50µl of OptiMEM (Gibco), incubated 10min and added to Huh-7.5 cells in a 48-well plate that had been seeded the day before with 3 x 10^4^ cells/well.

During the incubation period, the medium was changed to DMEM containing 1.5% FBS and 1% NEAA before transfection. To enhance transfection efficiency, cells were then subjected to a spinoculation process for 30min at 37°C and 1000g. Four hours post-incubation, the cells were lysed, and luciferase measurements were conducted using the Dual Luciferase Assay (Promega). The following settings were employed for the luciferase measurements: 10µl of cell lysate was mixed with 50µl of LARII, shaken for 2 sec, FLuc measurement was done for 5sec before injecting 50µl of Stop and Glo Reagent, and after 2sec of shaking, RLuc measurement was performed for 5sec. Luminescence was read on a FLUOstar Omega Microplate Reader with injectors (BMG Labtech).

### Full-genome HCV RNA transfection in Huh-7.5 cells

For assays with full-genome HCV RNAs, 100 ng of HCV chimeric RNAs were co-transfected with an equimolar amount of a capped monocistronic FLuc vector (FLuc:DEST40, kindly provided by Charles Rice, The Rockefeller University). This was done to normalize the IRES-driven GLuc HCV activity by the 5′ cap-driven FLuc activity. The remainder of the transfection protocol was consistent with the protocol previously described for bicistronic RNAs. Notwithstanding the secretion of GLuc, prior results indicated that at 4 hours post-transfection, the relative light units detected intracellularly exceed those detected extracellularly (Fig. S10). This observation informed the selection of this time point for the assays. Luciferase measurements were conducted in accordance with established protocols, with the exception that in this instance, GLuc activity was detected using the Stop and Glo Reagent. Luminescence was subsequently measured using a LUMIstar OPTIMA microplate reader (BMG Labtech). To assess the impact of IRES population heterogeneity on translational efficiency, the four IRES populations selected (Gt1a - samples 624 and 996, and Gt3a - samples 005 and 1017) were mimicked according to the variant frequency as determined by cloning methods. Consequently, 100ng of each population was comprised of a mixture of variants present in varying RNA amounts. The remaining transfection and luciferase measurement protocols were executed in strict accordance with the protocols outlined for full-length constructs. Experiments were done at least in triplicate with 3 technical replicates each.

### Statistical analysis

Statistical analyses were performed in Prism (GraphPad Prism Version 10.3.1 for Windows, GraphPad Software, Boston, Massachusetts, USA, www.graphpad.com). The statistical tests utilized, the number of biological replicates, as well as Fs, t-values, and degrees of freedom (df) are specified in each figure legend. In all cases, Shapiro-Wilk tests were employed to assess normality of the data. Statistical significance was defined as a *P* value ≤ 0.05.

### Mathematical model

We modelled cellular cofactors (F) (e.g., IRES binding factors such as ITAFs or translation initiation factors) using a logistic function, meaning that the concentration of F is allowed to increase up until a maximal concentration (K_F_). Similarly, multiple IRES variants (Ii) can coexist. IRES variants interact with cofactors with probability ***μ***, triggering the translation of viral proteins at a rate ***κ***. We constructed and compared three alternative models (Fig. S7). In model 1, there is no interaction between IRES variants, or cofactors and viral proteins (Fig. S7a). Model 2 introduces pairwise interactions between IRES variants, expressed as interaction or competition coefficients *α*ij. Such interactions affect the concentration of available IRES variants or their recycling (Fig. S7b). Model 3 allows, in addition to IRES interaction, the feedback between protein translation and the cofactors, with a coefficient *λ* (Fig. S7c). This model allows us to study the total change in protein yield (ΔPT) predicted by either the IRES interaction (model 2) or the IRES and protein-cofactors interaction (model 3), with respect to a no-interaction scenario (model 1). At equilibrium, we find:

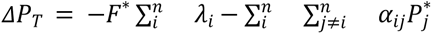

where F* is the concentration of cofactors and P* that of the proteins resulting from the n interacting IRES variants in the sample. This result implies that either the interaction between IRES variants (*α* ≠ 0, *λ* =0) or the interaction between cofactors and protein yield (*α* ≠ 0, *λ* ≠ 0) can independently and jointly, lead to increased protein translation provided that the competition coefficients defining those interactions, are negative (*α* < 0, *λ* < 0). In the context of viral protein translation, negative coefficients mean that the interaction between IRES variants can promote each other’s abundances or activities, their persistence or recycling, leading to larger levels of protein translation. Similar scenarios have mechanistic precedents in the literature(66, 68, 69) (A full description of the models and simulations is provided as part of the supplementary material).

## Data Availability Statement

The sequences in this study are available in the GenBank database (IRES master sequences—accession numbers PP350539 - PP350580, Sample 05 IRES clones— accession numbers PP350581 - PP350634, Sample 1021 IRES clones—accession numbers PP350635 - PP350688, Sample 624 IRES clones – accession numbers PV847832 - PV847922, Sample 996 IRES clones – accession numbers PV847923 - PV847977, Sample 1017 IRES clones – accession numbers PV847978 - PV848031, Sample 1016 IRES clones – accession numbers PV882825 - PV882903). NGS data are available under Bioproject number <u>PRJNA1078827</u>, and Biosamples <u>SAMN49662994</u> (996), <u>SAMN49662995</u> (624), <u>SAMN49662996</u> (1016), <u>SAMN40019051</u> (1021) and <u>SAMN40019050</u> (05).

## Code availability

Code to replicate supplementary figures 8 and 9 is available at the following link: https://github.com/eferrada/HCV_ProteinTranslation Shannon entropy for NGS data was calculated using custom scripts developed in Python for each position and normalized by dividing by the logarithm of the read count at that position, the script is available at the following link: https://github.com/NEcheverria1985/Shannon-entropy-from-NGS

## Acknowledgments

We would like to thank Dr. Charles Rice (Rockefeller University, New York, USA) and Dr. Marcelo López-Lastra (Pontificia Universidad Católica de Chile, Santiago, Chile) for kindly providing cells, vectors, and molecular clones for this work. In addition, we thank Stéphanie Beaucourt for technical assistance. This research was supported by Programa de Desarrollo de Ciencias Básicas (PEDECIBA— research grant Despegue Científico 2023-2024 to N.E.); Comisión Sectorial de Investigación Científica, Universidad de la República, Uruguay (CSIC—2018 R&D grant ID 288 to N.E.); Agencia Nacional de Investigación e Innovación (ANII—FMV_1_2014_1_104171 grant to P.M. and Mobility and training scholarship MOV_CA_2015_1_107441 awarded to N.E.); Comisión Académica de Posgrados (CAP, UdelaR, PhD Scholarship to N.E.); FOCEM (Fondo para la convergencia estructural del MERCOSUR–COF 03/11 to P.M and G.M.); and G4 Institut Pasteur de Montevideo program. The funders had no role in study design, data collection and analysis, decision to publish, or preparation of the manuscript.

## Authors’ Contributions

P.M., G.M., and N.E. conceived and designed the experiments, planned and supervised the project; N.E., P.P., F.G., and L.A. performed the experiments; N.E., M.S., and A.F. analyzed the data; N.H. collected the samples; E.F. performed the mathematical model. N.E., E.F., A.F., P.M., and G.M. wrote the original draft, and J.C. revised the final version. All authors have read and agreed to the published version of the manuscript.

## Materials & Correspondence

Requests should be addressed to Pilar Moreno and Gonzalo Moratorio.

## SUPPLEMENTARY MATERIAL

**Supplementary Table S1.** IRES mutant variants found as consensus sequences among Uruguayan HCV-chronically infected patients.

| HCV<br>Genotype | IRES<br>variant | Mutations <sup>a</sup> | Position within<br>secondary structure <sup>b</sup> | Frequency<br>(n) | Sample ID |
| --- | --- | --- | --- | --- | --- |
| 1a | HCV-1a | No changes | - | 2 | 28, 955 |
|  | 1 | G107A | D.II | 1 | 02 |
|  | 2 | A204C | subD.IIIb | 8 | 06, 07, 12, 18,<br>20, 21, 30, 1011 |
|  | 3 | A204U | subD.IIIb | 1 | 990 |
|  | 4 | Ins205U | subD.IIIb | 1 | 03 |
|  | 5 | A243G | D.III | 1 | 996 |
|  | 6 | G107A+A204C | D.II & subD.IIIb | 3 | 27, 598, 1020 |
|  | 7 | G107A+Ins207A | D.II & subD.IIIb | 1 | 08 |
|  | 8 | A204U+C340U | subD.IIIb & D.IV | 1 | 624 |
|  | 9 | A204C+A243G | subD.IIIb & D.III | 2 | 31, 32 |
|  | 10 | G107A+A204C+A243G | D.II, subD.IIIb & D.III | 3 | 22, 25, 1019 |
|  | 11 | A119C+A204C+U248C | D.II, subD.IIIb & D.III | 1 | 1021 |
|  | 12 | A204C+U248C+A358U | subD.IIIb, D.III & core | 1 | 1018 |
| 1b | HCV-1b | No changes | - | 8 | 01, 10, 13, 14,<br>26, 34, 36, 39 |
|  | 13 | C204U | subD.IIIb | 2 | 16, 24 |
|  | 14 | Ins207A | subD.IIIb | 1 | 15 |
|  | 15 | Ins207A+A214U | subD.IIIb | 1 | 04 |
| 3a | HCV-3a | No changes | - | 0 | - |
|  | 16 | U247C | D.III | 1 | 991 |
|  | 17 | A119C+G203A | D.II & subD.IIIb | 1 | 1017 |
|  | 18 | A119C+G203A+G243A | D.II, subD.IIIb & D.III | 2 | 05, 982 |
|  | 19 | A119C+G203A+Ins207A+G243A | D.II, subD.IIIb & D.III | 1 | 1016 |

<sup>a</sup> According to reference strains H77 (1a, NCBI accession number: AF009606), Con1 (1b, NCBI accession number: AJ238799) & NZL1 (3a, NCBI accession number: D17763).
<sup>b</sup> **D:** Domain; **subD:** Subdomain

**Supplementary Figure S1.**
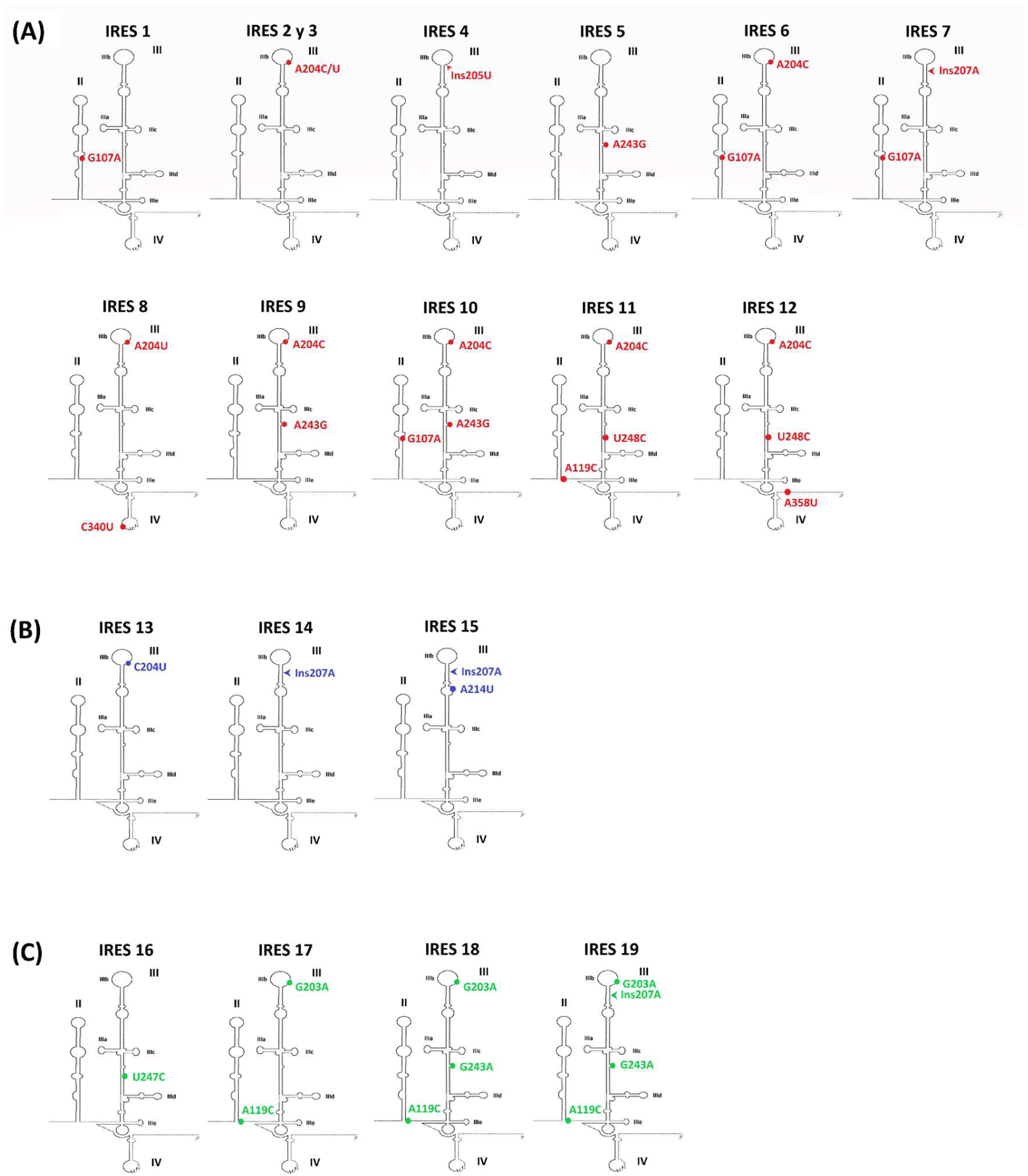
Location of the mutations found in the consensus IRES, whose translational efficiencies were evaluated. The secondary structure of the IRES (domains II to IV of the 5’NCR region) as proposed by Honda et al. (1999)(15) is schematized, and the 19 different mutation combinations evaluated at the translational level are indicated. Nucleotide changes are indicated by colored dots, while insertions are indicated by arrows. **A)** IRES of Gt1a, mutations in red. **B)** IRES of Gt1b, mutations in blue. **C)** IRES of Gt3a, mutations in green.

**Supplementary Figure S2.**
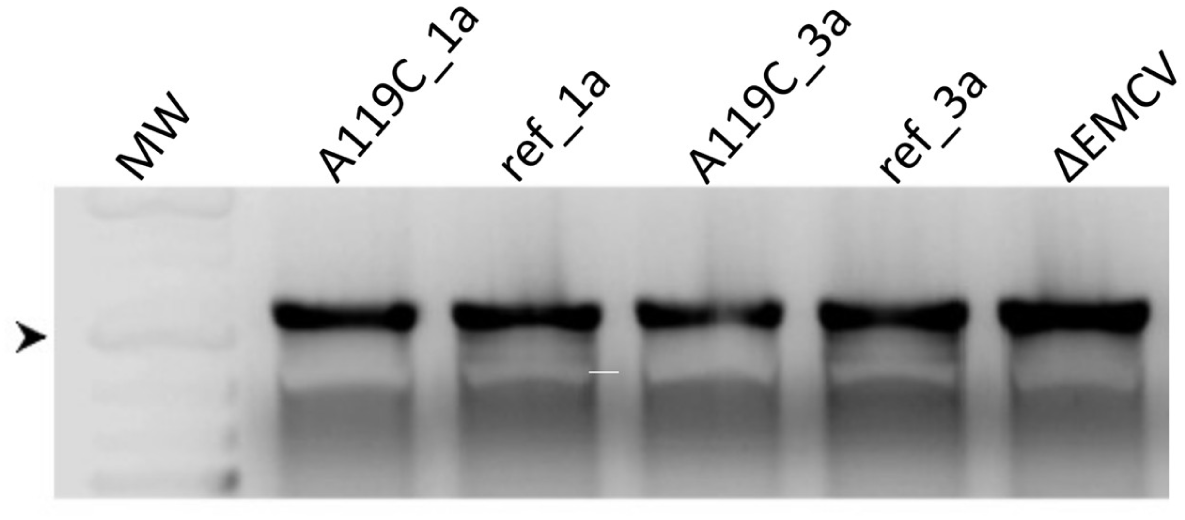
Non-denaturing agarose gel of bicistronic *in vitro* transcribed RNAs. 1% agarose native gel for visualization of bicistronic RNA integrity from a representative experiment. SYBR Safe was used as an intercalating agent. Lanes 2-3: GT1a IRES (mutant A119C and reference sequence for Gt1a). Lanes 4 and 5: Gt3a IRES (mutant A119C and reference sequence for Gt3a). MW: molecular weight of double-stranded DNA, 1 Kbp Plus (Fermentas). Lane 6: negative control (IRES DL_ΔEMCV). 1 µg of RNA was mixed with an equal volume of RNA Loading Dye 2X, pre- heated for 10 min at 70°C, and then loaded onto the gel. The arrowhead indicates the band corresponding to 1.5 Kbp. All *in vitro* transcribed RNAs are not degraded and seem to have the same apparent molecular weight.

**Supplementary Figure S3.**
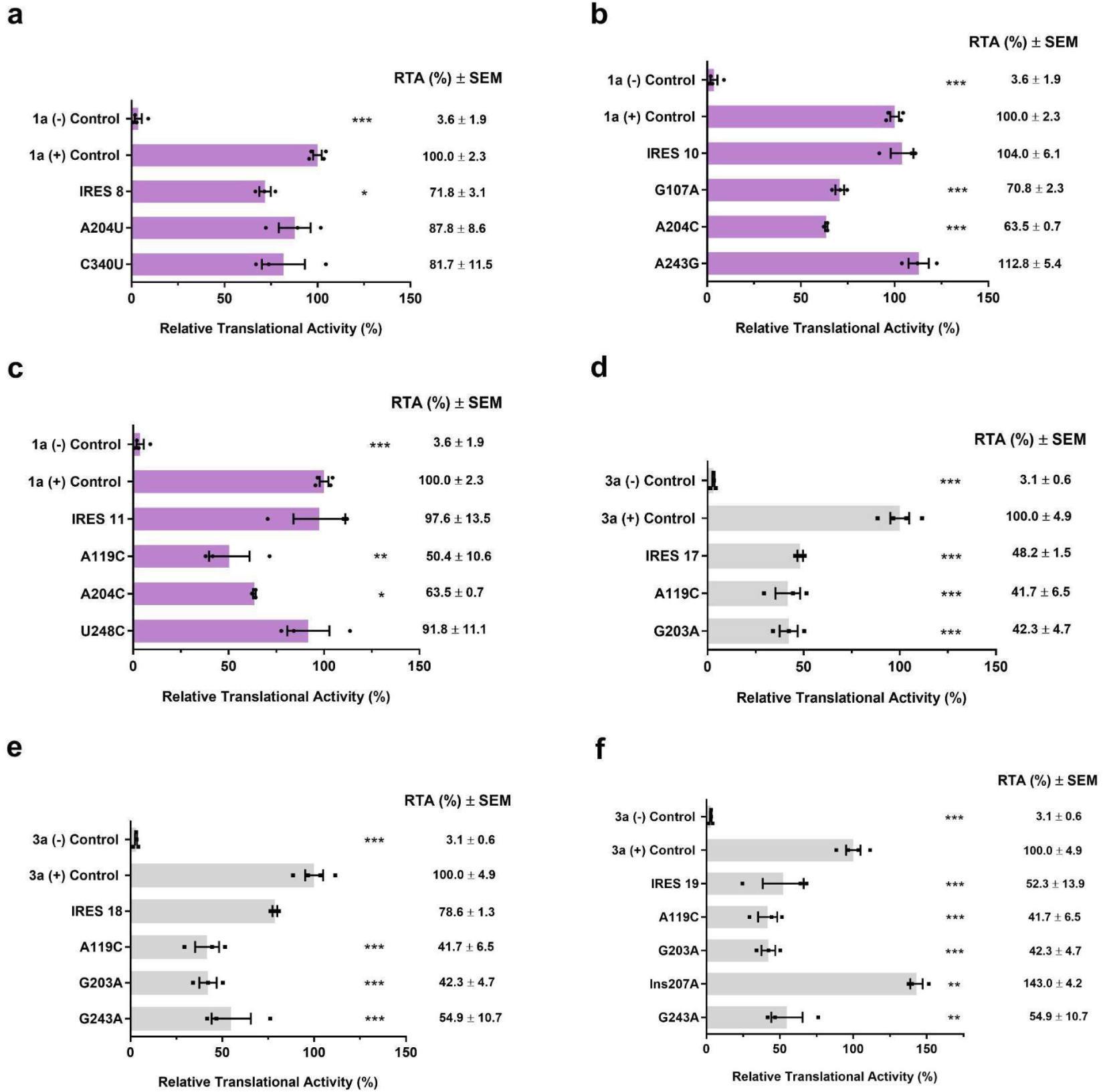
Dissecting the contribution of specific point mutations to *ex vivo* relative translation activity of master IRES variants harboring multiple mutations. Master IRES variants carrying multiple mutations are presented, along with their respective point single mutants. The controls are displayed at the top of each panel. **(A)**, **(B)**, **(C)** GT1a IRESs: IRES 8, 10, and 11, respectively (faded purple bars, black circles). **(D)**, **(E)**, **(F)** GT3a IRESs: IRES 17, 18, and 19, respectively (faded grey bars, black squares). The relative translational activity was evaluated by comparing it with its respective positive control (1a-Jc1-GLuc/GNN, or 3a-Jc1-GLuc/GNN), and mutants 1a-266+268-Jc1- GLuc/GNN, or 3a-266+268-Jc1-GLuc/GNN were employed as negative controls (inefficient). The firefly luciferase (FLuc) and Gaussia luciferase (GLuc) activities were quantified, with the GLuc/FLuc ratio serving as the activity index. The ratio corresponding to each of the positive controls was arbitrarily designated as 100%. The means and standard errors (SEM) of at least three independent experiments are plotted. Asterisks indicate IRESs with significant differences in their efficiency (one-way ANOVA followed by a Dunnett’s multiple comparison test: *** *p*<0.001, ** *p*<0.01, * *p*<0.05). **(A)** IRES 8: F-value = 47.95, df = 4; **(B)** IRES 10: F-value = 156.8, df = 5; **(C)** IRES 11: F-value = 27.72, df = 5; **(D)** IRES 17: F-value = 83.09, df = 4; **(E)** IRES 18: F-value = 43.87, df = 5; **(F)** IRES 19: F-value = 42.72, df = 6.

**Supplementary Table S2.** Samples selected for intra-host analyses.

| Genotype | Sample ID | Master IRES Nr | Mutations | *in vitro* RTA<br>± SEM (%) | *ex vivo* RTA<br>± SEM (%) | Mean RTA<br>difference (%) | P value |
| --- | --- | --- | --- | --- | --- | --- | --- |
| 1a | 996 | 5 | A243G | 68,5 ± 2,5 | 104,3 ± 9,4 | 35,8 | <0.001 |
|  | 624 | 8 | A204U+C340U | 95,9 ± 3,8 | 75,9 ± 3,7 | -20,0 | >0.05 |
|  | 1021 | 11 | A119C+A204C+U248C | 80,3 ± 4,0 | 97,6 ± 13,5 | 17,2 | >0.05 |
| 3a | 1017 | 17 | A119C+G203A | 32,5 ± 2,3 | 60,8 ± 9,6 | 28,3 | <0.001 |
|  | 05 | 18 | A119C+G203A+G243A | 20,2 ± 2,0 | 76,7 ± 4,5 | 56,5 | <0.001 |
|  | 1016 | 19 | A119C+G203A+Ins207A+<br>G243A | 28,1 ± 0,4 | 52,3 ± 13,9 | 24,2 | >0.05 |

**Supplementary Figure S4.**
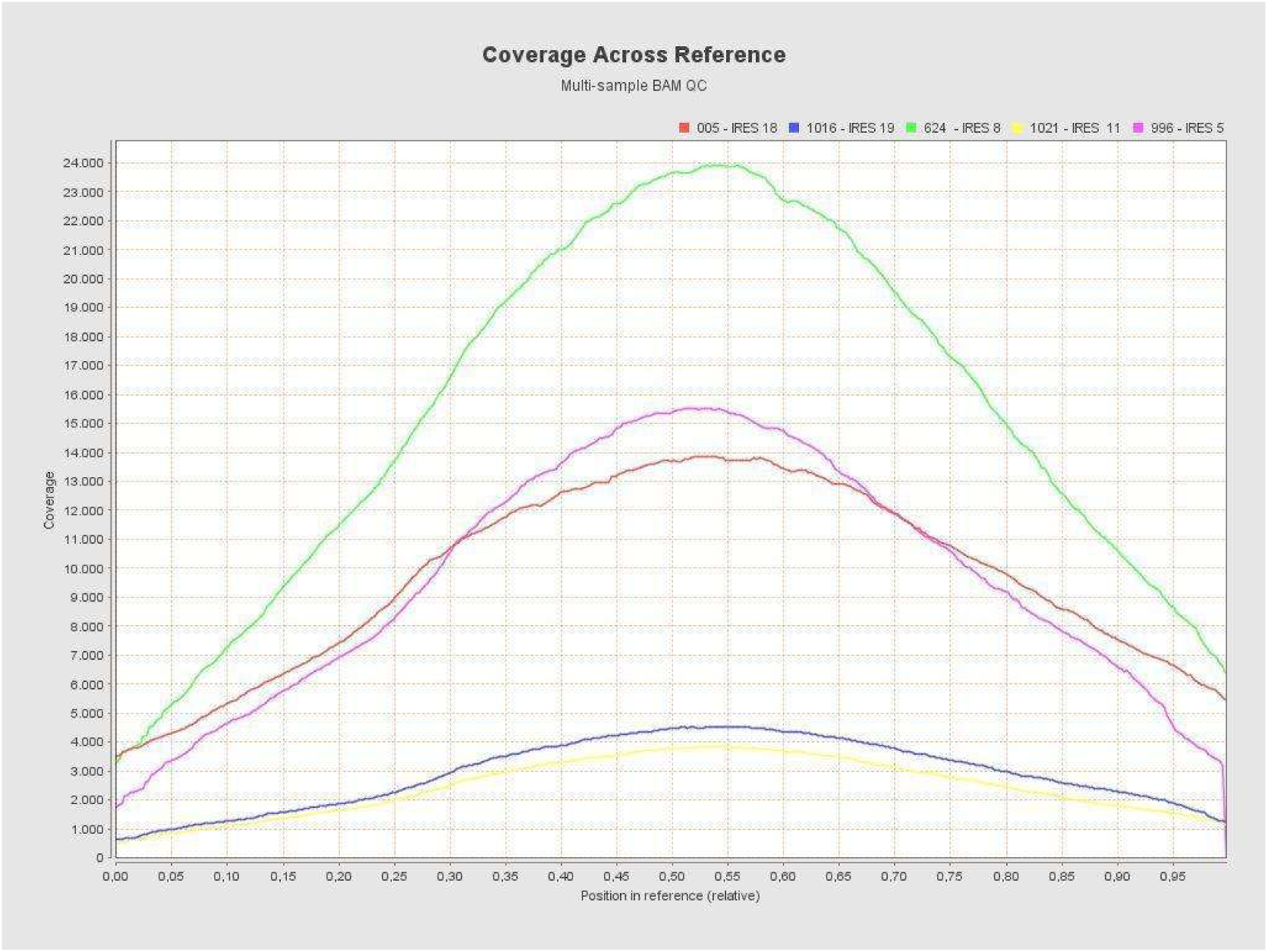
Coverage along the IRES following the alignment of the filtered reads. The different samples and their corresponding IRES are indicated in different colors. The graph was generated with the Qualimap v2.3 tool(82).

**Supplementary Table S3.** Sample statistics obtained from multi-sample BAM QC analysis generated by Qualimap v.2.3 tool. (**82**).

| Sample ID -<br>IRES Nr | Coverage<br>mean | Coverage<br>std* | GC<br>% | Mapping<br>quality<br>mean | Insert<br>size median |
| --- | --- | --- | --- | --- | --- |
| 996 - IRES 5 | 9588 | 4156 | 59.19 | 59.97 | 190 |
| 624 - IRES 8 | 15352 | 6201 | 58.75 | 59.99 | 162 |
| 1021 - IRES 11 | 2431 | 983 | 59.72 | 59.98 | 180 |
| 005 - IRES 18 | 9704 | 3139 | 59.91 | 59.98 | 192 |
| 1016 - IRES 19 | 2893 | 1179 | 58.71 | 59.97 | 193 |

\* std - Standard deviation

**Supplementary Figure S5.**
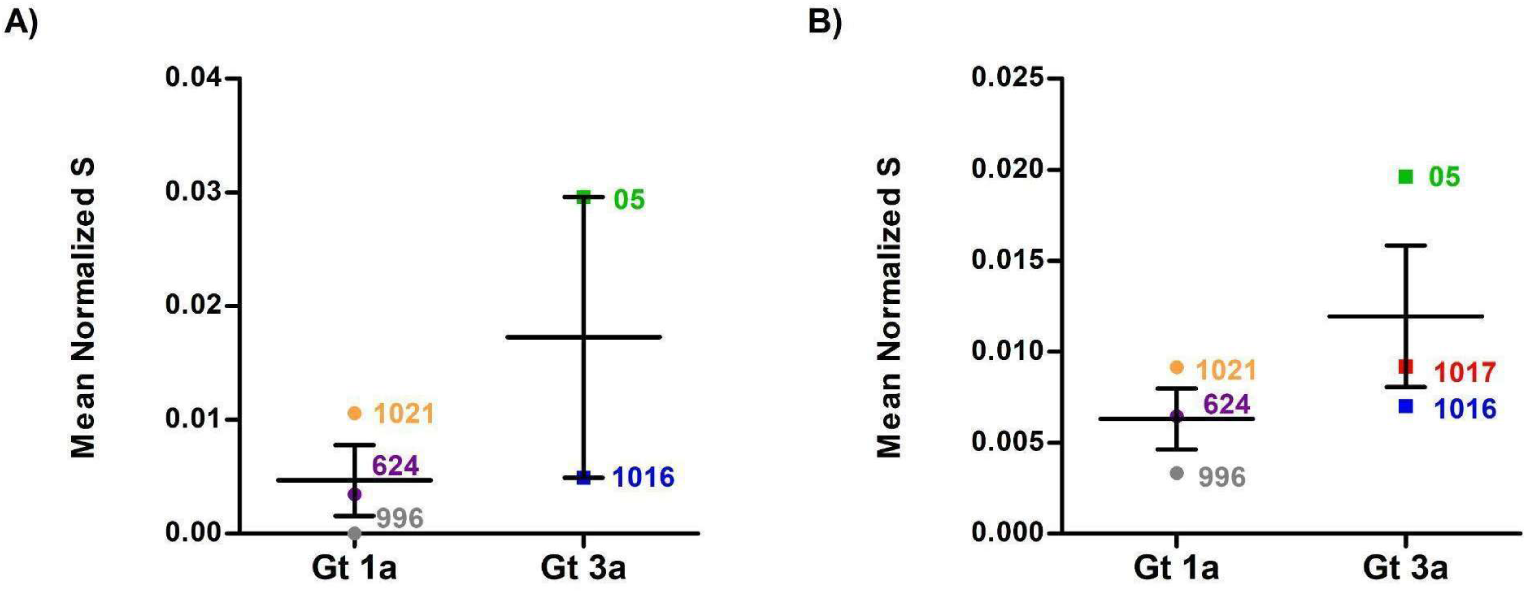
Mean Normalized Shannon entropy per sample according to genotype. Each dot/square represents the mean normalized entropy of each sample (as plotted in Figure 5). The mean and standard errors (SEM) of the entropy values for each genotype are indicated. The colored dots represent samples corresponding to Gt1a (grey: 996, violet: 624, orange: 1021), whereas the colored squares represent those corresponding to Gt3a (red: 1017, green: 05, blue: 1016). **A)** Results of NGS data; **B)** Results of IRES cloning.

**Supplementary Table S4.** Results from the molecular cloning approach of IRES of interest.

| Sample ID | Master IRES Nr | Clones (n) | IRES haplotypes (n) | Master Sequence | Master Sequence % (n) | Mixed Infection | Major genotype | Minor genotype | Minor genotype % (n) |
| --- | --- | --- | --- | --- | --- | --- | --- | --- | --- |
| 996 | 5 | 55 | 11 | A243G | 80,00 (44) | No | - | - | - |
| 624 | 8 | 91 | 20 | A204U+C340U | 70,33 (64) | Yes | 1a | 3a | 1,10 (1) |
| 1021 | 11 | 54 | 20 | A119C+A204C+U248C | 42,59 (23) | Yes | 1a | 3a | 1,85 (1) |
| 1017 | 17 | 54 | 14 | A119C+G243A | 75,93 (41) | Yes | 3a | 1a | 1,85 (1) |
| 05 | 18 | 54 | 23 | A119C+G203A+G243A | 51,85 (28) | Yes | 3a | 1a - 1a/3a* | 7,41 (4) |
| 1016 | 19 | 79 | 35 | A119C+G203A+Ins207A+G243A | 34,18 (27) | No | - | - | - |

\* Recombinant sequences(45).

**Supplementary Table S5.**
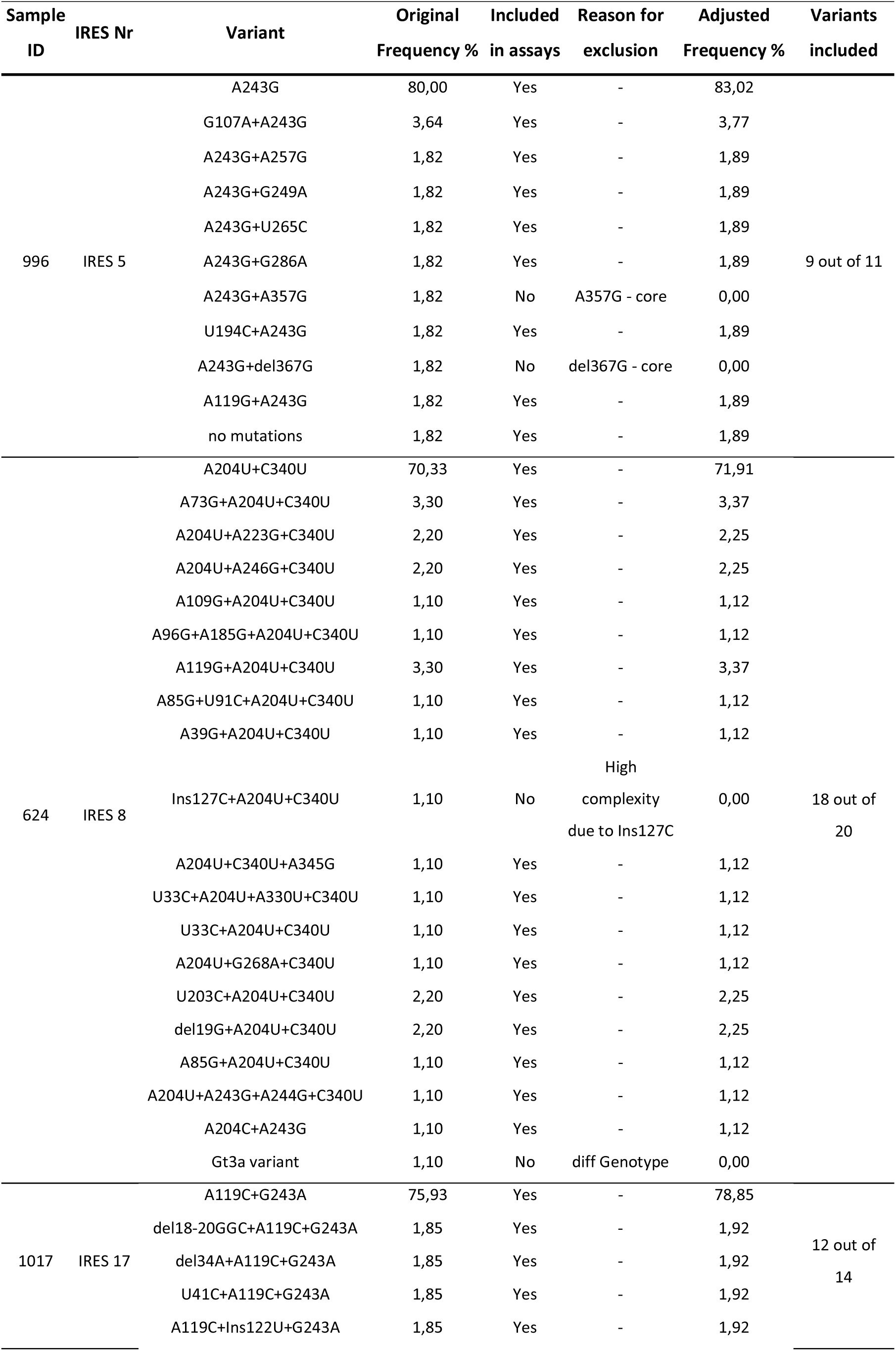
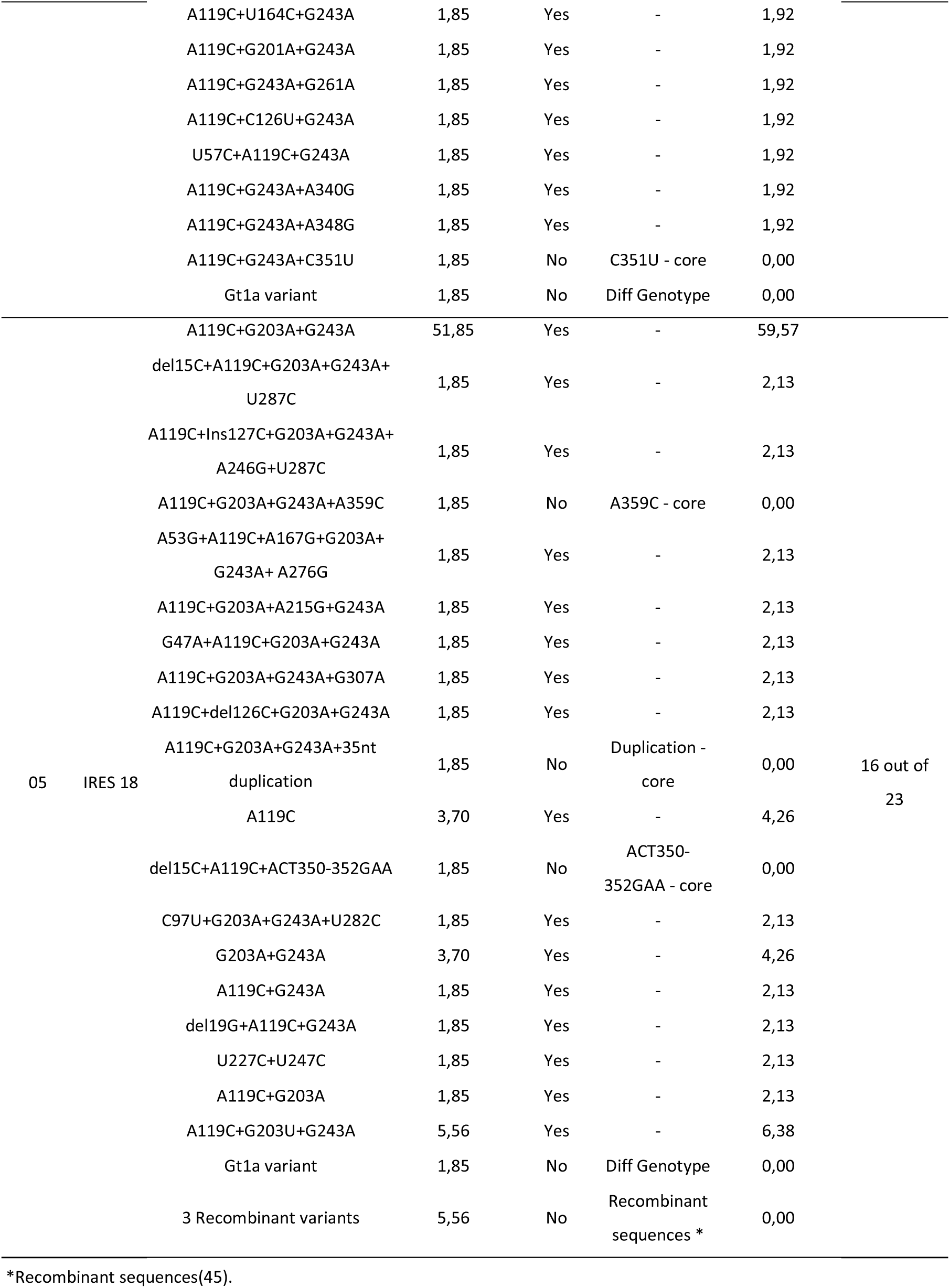
Samples selected for analysis of IRES heterogeneity and their corresponding variants.

**Supplementary Figure S6.**
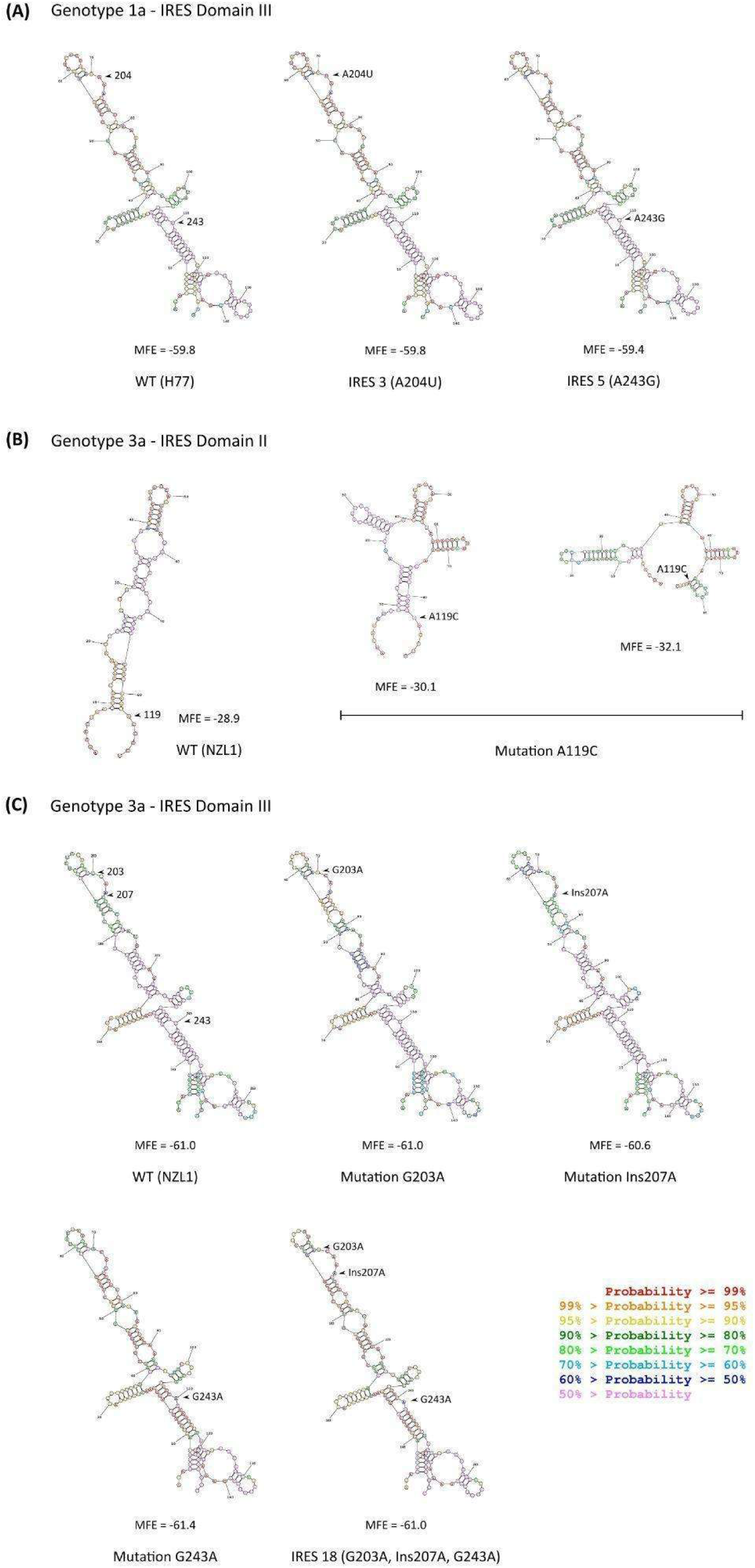
Predicted secondary structures of IRES domains II and III of less efficient HCV mutants. Secondary structures were generated with RNAstructure Webserver, version 6.5 (available at https://rna.urmc.rochester.edu/RNAstructureWeb/). The secondary structures of the reference sequences are displayed in the left panels for genotypes 1a - domain III (A), 3a - domains II (B), and III (C). The different mutants of interest are schematized in the right and lower panels. Base pairing probabilities are indicated by different colors, as shown in the bottom right corner of the figure. The minimum free energy of the structures is detailed below each. Domain II encompasses nucleotides 36 to 124, while domain III spans nucleotides 134 to 290.

**Supplementary Figure S7.**
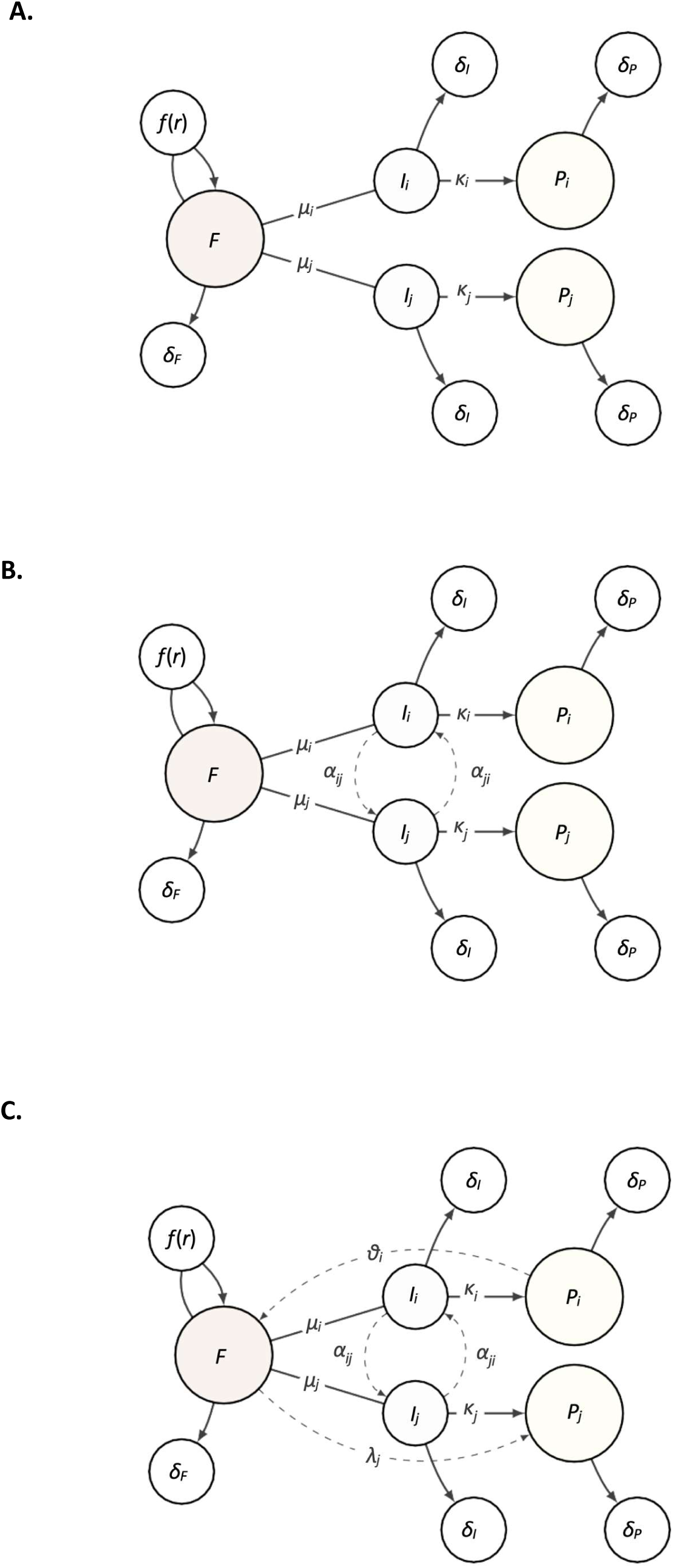
**Graphical description of three alternative models for IRES-mediated translation dynamics in HVC. *A.*** Model 1 does not assume interactions between IRES variants or between cofactors and protein yield. In ***B.*** Model 2, IRES variants are allowed to interact, affecting each other abundances and, as a consequence, protein translation. ***C.*** Model 3, assumes both interacting IRES variants, and feedback between protein yield and cofactors. For details about the models, see main text.

### Supplementary Data

#### Overall description and Model 1

These models describe alternative scenarios for the translation dynamics of hepatitis virus C (HCV) proteins mediated by IRES. It is similar to a resource- consumer model of species interaction in which the replication of variants of the virus uses common cofactors of resources available in the cell, and where variant species might eventually interact competitively or cooperatively.

*F* stands for cofactors or cellular resources (eg IRES binding cofactors, amino acids, ribosomes, etc.) that are essential for IRES-mediated translation and might vanish after persistent viral replication. Cofactors concentration decays at a rate *δ_F_* and has a resource renewal function *f* (*r*), which we describe as a logistic function. In the case of Model 1 (Figure S7A), cofactors are not influenced by the concentration of IRES or viral proteins alone. For this model, the change of *F* over time can be expressed as:

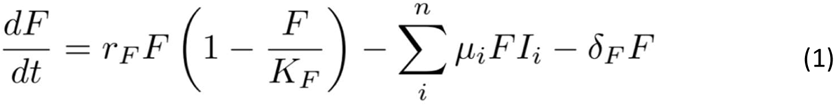

where *r_F_* is the overall production rate of cofactors, and *K_F_*, their maximal concentration in the cell. Similarly, the overall change in the concentration of IRES variants (*Ii*) is:

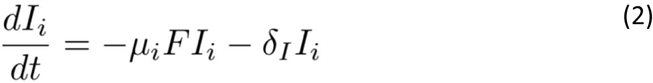

Initially, IRES *Ii* has an available concentration which declines as it binds co-factors with probability *µ*. We assume that IRES variants, *Ii*, decay at a rate *δI*, proportionally to their available concentration.

Lastly, for Model 1, the overall change in the concentration of proteins (*Pi*) is proportional to the cofactor-bound IRES, at a rate *κi*, and can be expressed as:

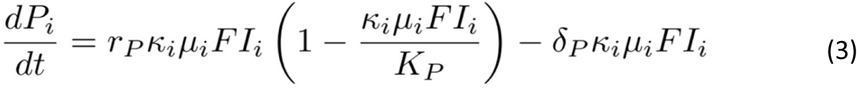

where *r*P is the overall rate of protein yield and *KP*, the maximum concentration of protein in the medium. We might consider the total protein concentration (*PT*), which at equilibrium can be written as:

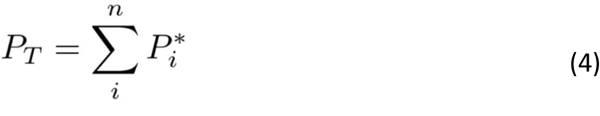

Because there is no feedback between IRES variants or between co-factors and protein concentration, we call Model 1, the no-interaction model, and it can be summarized as:

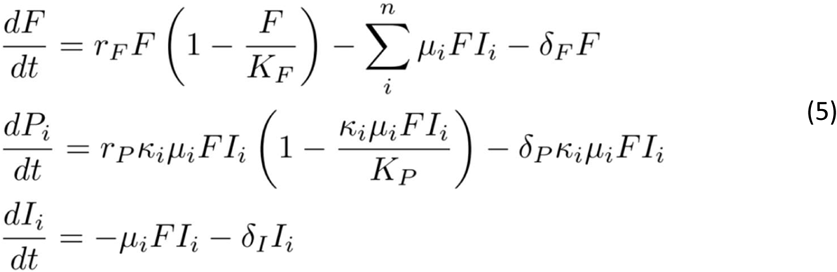

Without loss of generality, we assume *δP* and *δI* rates are the same for all *i*.

#### Model 2: IRES interaction model

In an alternative model, IRES variants are allowed to interact with each other, ultimately impacting viral protein yield (Figure S7B). IRES interactions might arise from several mechanisms, such as through ribonucleoproteins, through trans-complementation (for details, see main text). The model can be summarized as:

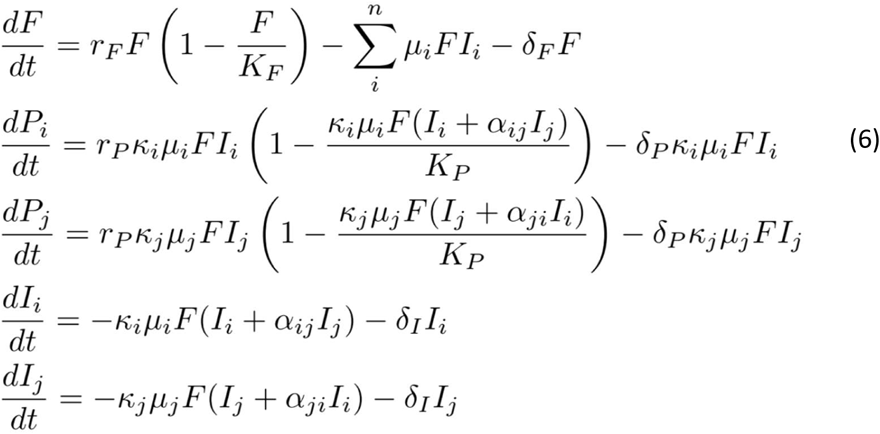

coefficients *α* define the impact of IRES variants into each other’s abundances. These coefficients can also be interpreted as affecting the general availability or recycling of IRES variants. Generally, there will be *n*(*n* − 1) *α* coefficients, or pairwise interactions between IRES variants.

#### Model 3: Protein-cofactors and IRES-IRES interaction model

We consider yet an additional model in which both IRES variants, as well as proteins and co-factors are allowed to interact (Figure S7C). Protein yield, for instance, might induce the synthesis of more cofactors or limit their overall concentration. We introduce two additional coefficients (*θ* and *λ*), assuming that the interaction between protein yield and co-factors is asymmetric. Such model can be summarized as:

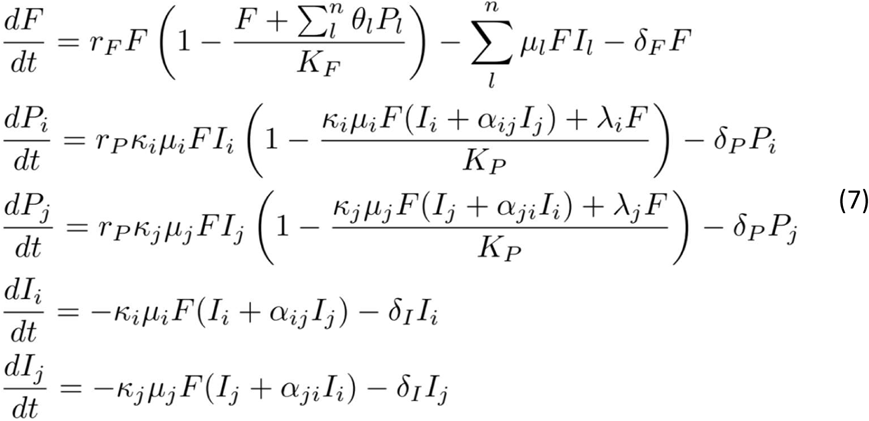

If these coefficients (*α*, *λ* and *θ*) are set 0, we recover Model 1 (*α* = *λ* = *θ* = 0) and Model 2 (*α* = 0; *λ* = *θ* = 0). In the general context of the resource-consumer model, the values of these coefficients define the interaction dynamics between species, leading to alternative competition scenarios.

#### Analyses of equilibria

We study the more general model, Eq (7), with both IRES and protein-co-factors interactions. At equilibrium for *dF/dt* = 0, we have:

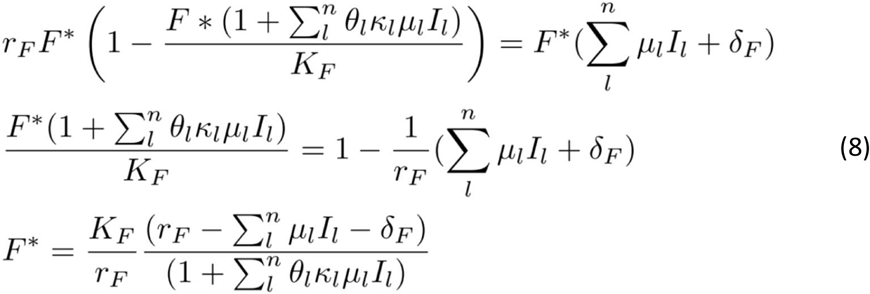

Because *I* should be very small at equilibrium, we have:

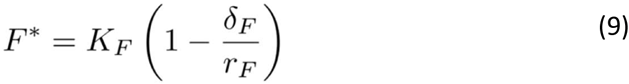

Because the concentrations of cofactors cannot be lower than zero, *r_F_ > δ_F_ >* 0; and because Eq. (9) is independent of competition coefficients (i.e., *α*, *λ*, and *θ*), this result applies to all three models.

Similarly, at equilibrium for *Pi*, we have:

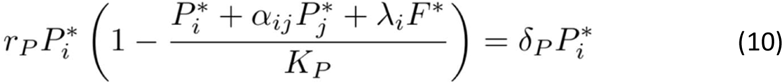

which, for a general model with *n* IRES variants, it gives:

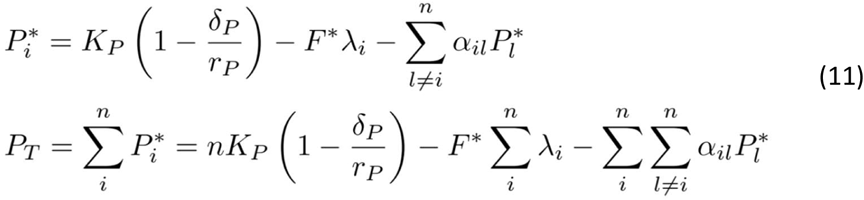

Interestingly, Eq. (11) is independent of the coefficient *θ*, which suggests that the interaction between proteins and cofactors is asymmetric, and only the effect of cofactors on protein concentration would have a significant effect. The reason is that in the model, cofactor concentrations are assumed to be generally in excess with respect to the concentration of proteins, which depend on the intermediary role of IRES variants.

By comparing the total protein yield between Model 1 versus Models 2 and 3, and using Eq. (9) and (11), we get:

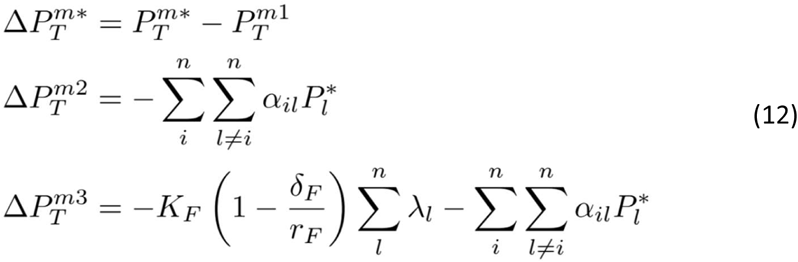

Because here we have *r_F_ > δ_F_ >* 0 and both *K_F_* and *Pi >* 0; Eq. (12) implies that for models 2 and 3 to lead to larger protein translation yields with respect to Model 1, competition coefficients *α* and *λ* should be negative.

In the case of Model 2, which describes the interactions between the IRES variants, competition coefficients *α <* 0 would lead to larger protein yields. The model also suggests that as long as there is a single dominant negative coefficient, protein yield can still be larger than in Model 1. In the context of resource-consumer models in ecology, negative *α* coefficients are associated with mutualistic interactions, compared to other combinations of coefficients that lead to competitive or parasitic scenarios. Similarly, and provided *λ <* 0, Model 3, which introduces interactions between proteins with resource cofactors, should lead to even larger protein yields. The model also predicts that even in the absence of interactions between IRES variants, the interaction between co-factors and proteins alone, could lead to increased protein yields.

Figure S8 shows numerical simulations contrasting results of Model 1 and Model 2, at a positive (Figure S8A-B) and negative (Figure S8C-D) values of the *α* parameter. As predicted by Eq. (12), at *αij* = *αji <* 0, Δ*P^m^*^2−*m*1^ *>* 0.0. Similarly, Figure S9 presents simulations comparing Model 1 and Model 3, with positive (Figure S9A-B) and negative (Figure S9C-D) coefficients (*α* and *λ*). As predicted by Eq. (12), at *αij* = *αji <* 0 and *λ <* 0, Δ*P^m^*^3−*m*1^ *>* 0.0.

**Supplementary Figure S8.**
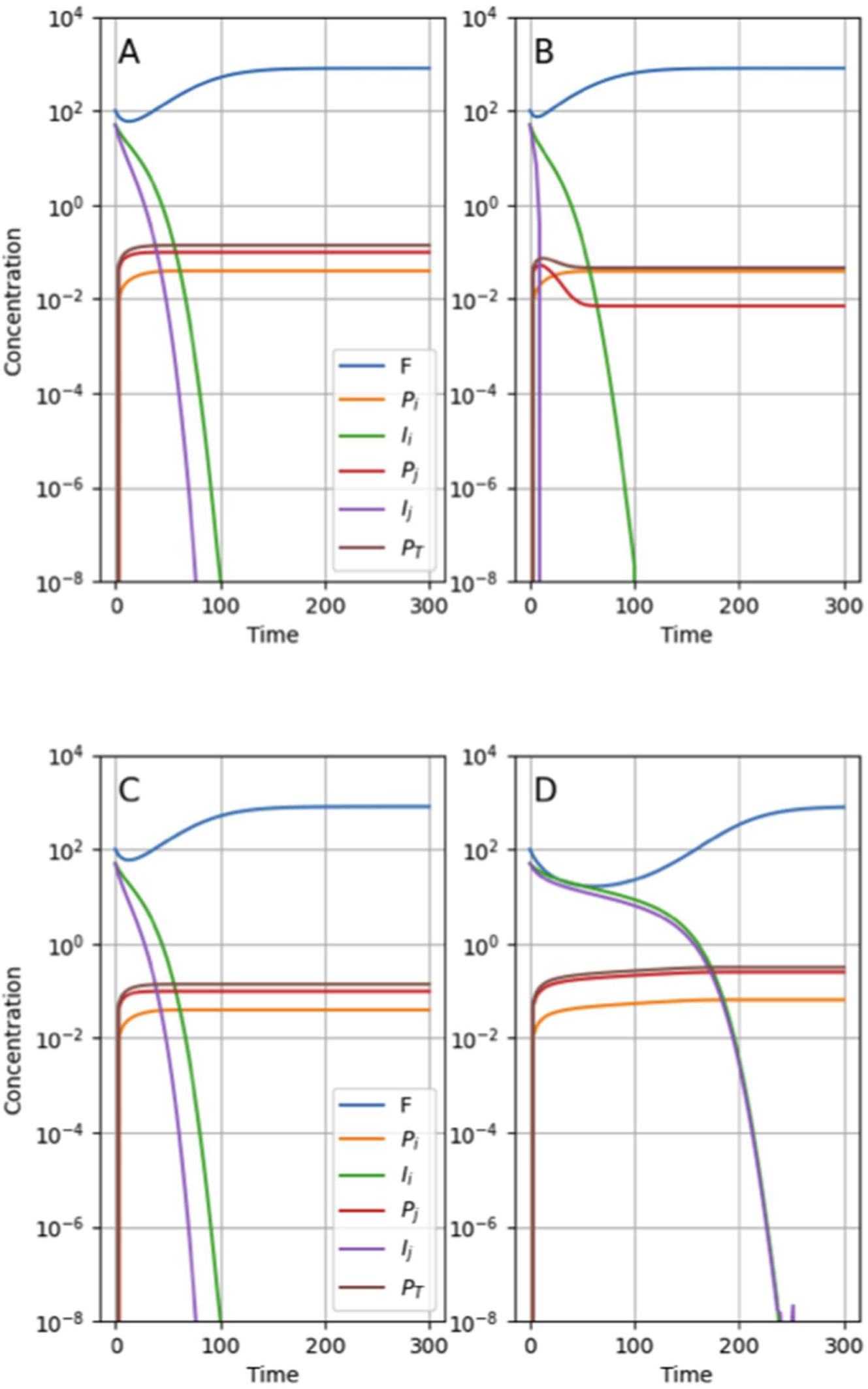
Numerical simulation comparing the no-interaction ( Model 1, A and C) and the IRES variants interaction (Model 2, B and. **D)**. We simulated Model 1 and Model 2, as described in Eqs. (5) and (6), respectively. We used 2 IRES variants, and for each scenario we followed the changes in concentration over t=300 iterations. Parameters were set as: *r_F_* = *rP* = 0.05; *K_F_* = *KP* = 1000; *δ_F_* = *δP* = 0.01; *δI* = 0.01; *µi* = 0.0011; *µj* = 0.0021; *κi* = 0.02; *κj* = 0.05. Competitions coefficients were set as (**A** and **B**) *αij* = *αji* = 0.5; and (**C**,**D**) *αij* = *αji* = −0.5.

**Supplementary Figure S9.**
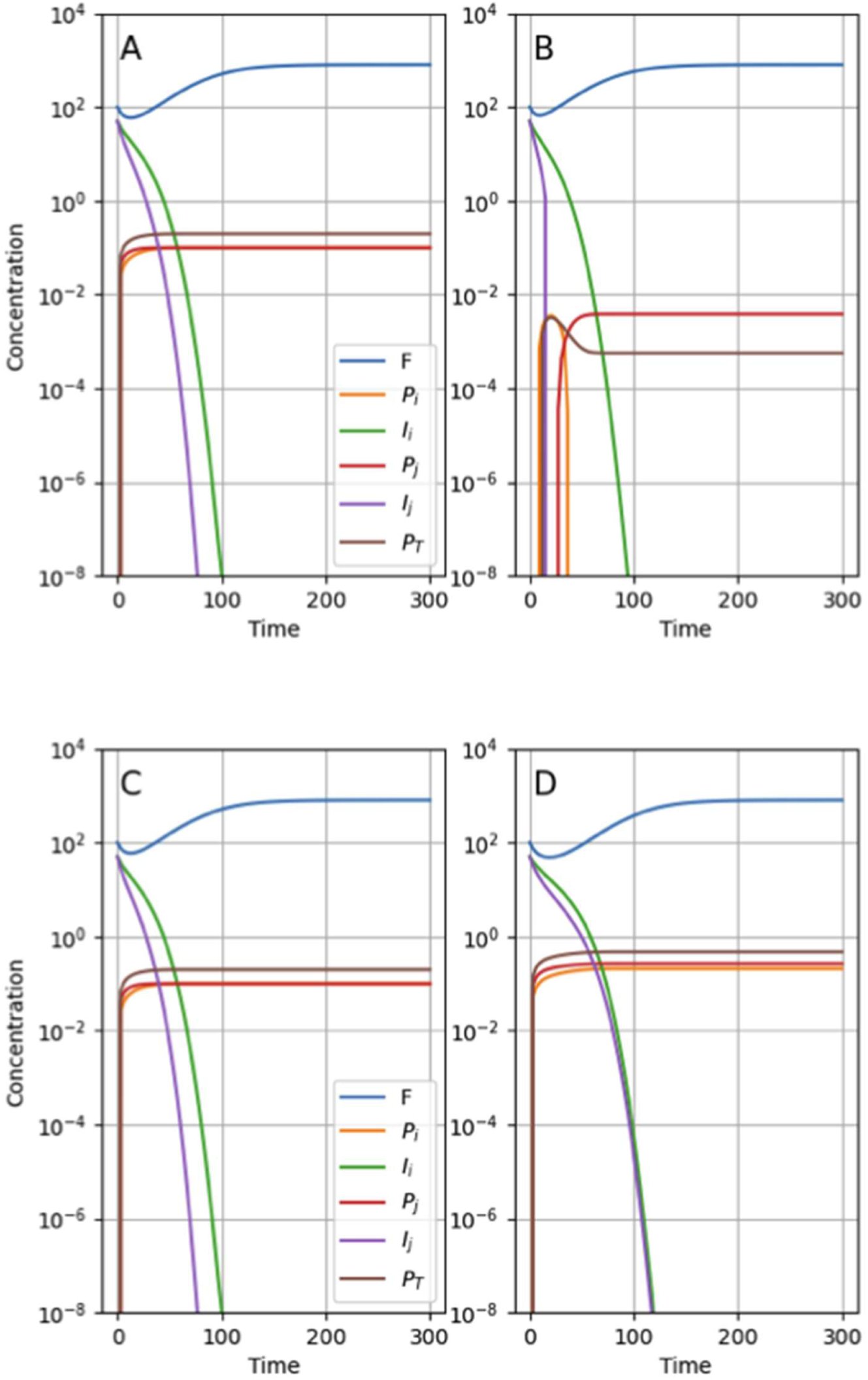
Numerical simulation comparing the no-interaction (Model 1, A and C) and the model with both cofactors and IRES variants interaction (Model 3, B and D). We simulated Model 1 and Model 3, as described in Eqs. (5) and (7), respectively. We used 2 IRES variants, and for each scenario we followed the changes in concentration over t=300 iterations. Parameters were set as: *r_F_* = *rP* = 0.05; *K_F_* = *KP* = 1000; *δ_F_* = *δP* = 0.01; *δI* = 0.001; *µi* = 0.0011; *µj* = 0.0021; *κi* = 0.02; *κj* = 0.05. Competitions coefficients were set as (**A** and **B**) *αij* = *αji* = 0.2; *θi* = *θj* = 0.0; *λi* = *λj* = 10 and (**C**,**D**) *αij* = *αji* = −0.2; *θi* = *θj* = 0.0; *λi* = *λj* = −10.

**Supplementary Table S6.** Demographic characteristics of patients enrolled in this study, virological parameters, and HCV genotype according to NS5B phylogeny.

| Sample | Sex | Age<br>(years) | Viral Load<br>(log10) <sup>a</sup> | Antiviral<br>therapy <sup>b</sup> | Response to<br>treatment <sup>c</sup> | NS5B<br>genotype |
| --- | --- | --- | --- | --- | --- | --- |
| 598 | M | 53 | 5.78 | PR | NR | 1a |
| 624 | M | 21 | 6.80 | ND | ND | 1a |
| 955 | F | 65 | 6.61 | PR | ND | 1a |
| 982 | M | 45 | 5.08 | PR | NA <sup>d</sup> | 3a |
| 990 | M | 40 | 6.71 | PR | SVR | 1a |
| 991 | M | 48 | 7.16 | ND | ND | 3a |
| 996 | F | 72 | 5.62 | ND | ND | 1a |
| 1011 | M | 31 | 6.48 | ND | ND | 1a |
| 1016 | M | 46 | 6.09 | ND | ND | 3a |
| 1017 | M | 58 | 6.17 | *Naïve* | NA | 3a |
| 1018 | M | 35 | 6.31 | *Naïve* | NA | 1a |
| 1019 | M | 46 | 6.39 | *Naïve* | NA | 1a |
| 1020 | M | 40 | 6.08 | *Naïve* | NA | 1a |
| 1021 | M | 67 | 5.79 | ND | ND | 1a |
| 001 | M | 51 | 6.37 | PR-PRT | NR-NR | 1b |
| 002 | F | 53 | 3.75 | *Naïve* | NA | 1a |
| 003 | F | 45 | 4.30 | *Naïve* | NA | 1a |
| 004 | F | 71 | ND | *Naïve* | NA | 1b |
| 005 | F | 52 | 8.09 | *Naïve* | NA | 3a |
| 006 | M | 45 | 7.00 | PR | NR | 1a |
| 007 | F | 35 | 8.00 | *Naïve* | NA | 1a |
| 008 | F | 51 | ND | *Naïve* | NA | 1a |
| 010 | F | 34 | 4.85 | PR | NR | 1b |
| 012 | M | 45 | 6.65 | *Naïve* | NA | 1a |
| 013 | M | 64 | 4.70 | PR | NR | 1b |
| 014 | M | 51 | 5.39 | PR-PRT | NR-NR | 1b |
| 015 | M | 30 | ND | *Naïve* | NA | 1b |
| 016 | F | 63 | 6.29 | *Naïve* | NA | 1b |
| 018 | M | 47 | ND, 6.20 | PR,<br>SOF/LDV | NR, Relapse<br><sup>e</sup> | 1a |
| 020 | M | 40 | 6.00 | PR | NR | 1a |
| 021 | M | 32 | 6.01 | *Naïve* | NA | 1a |
| 022 | M | 49 | 6.24 | PR | NR | 1a |
| 024 | M | 36 | ND | *Naïve* | NA | 1b |
| 025 | M | 48 | 6.90 | PR | NR | 1a |
| 026 | M | 59 | ND | IR | NR | 1b |
| 027 | M | 54 | 6.30, 7.10 | PR,<br>SOF/LDV | NR, Relapse<br><sup>e</sup> | 1a |
| 028 | M | 42 | 4.36 | PR | NR | 1a |
| 030 | M | 37 | 5.30 | *Naïve* | NA | 1a |
| 031 | M | 45 | ND | PR | NR | 1a |
| 032 | M | ND | 5.10 | *Naïve* | NA | 1a |
| 034 | F | 58 | 5.60 | PR | NR | 1b |
| 036 | F | 54 | 5.30 | *Naïve* | NA | 1b |
| 039 | M | 43 | 6.49 | PR | NR | 1b |

<sup>a</sup> Viral load at the time of sample collection. Log<sub>10</sub> of viral load determined in international units per milliliter (IU/mL) with commercial kits COBAS®TaqMan HCV Test or Real Time HCV (Abbot), v2.0. <1.40 indicates detectable values, yet not quantifiable.
<sup>b</sup> **PR**: peg-IFN- $\alpha$ /ribavirin; **Naïve**: no treatment received; **PRT**: PR+telaprevir; **SOF/LDV**: sofosbuvir/ledipasvir; **IR**: IFN- $\alpha$ /ribavirin
<sup>c</sup> **NR**: Non-responder; **SVR**: Sustained virological response
<sup>d</sup> Treatment suspended due to intolerance to interferon
<sup>e</sup> 2 samples from the same patient were collected, after non-response to PR and after relapse to SOF/LDV
**ND**: Not determined or unknown
**NA**: Not applicable

**Supplementary Table S7.**
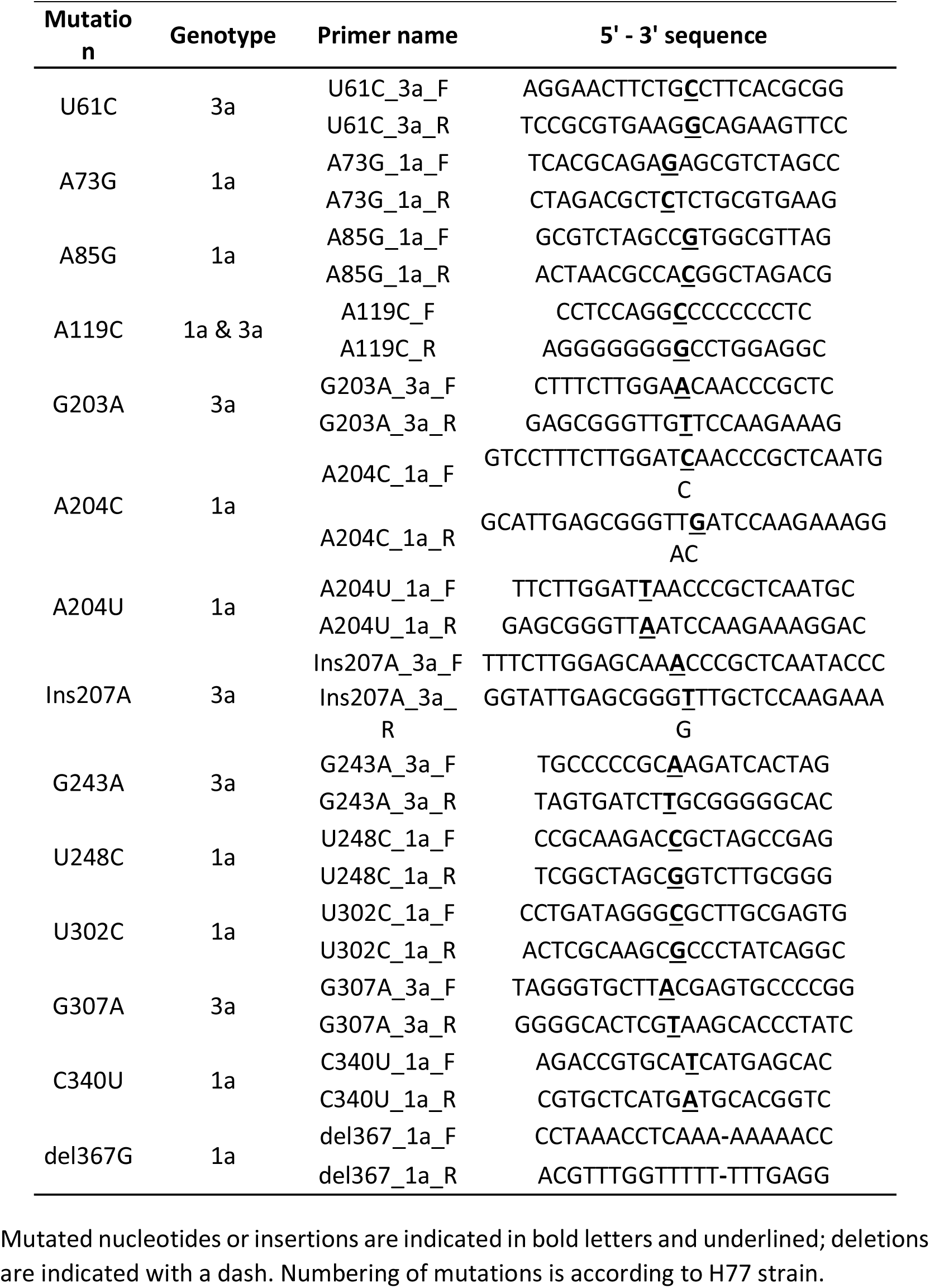
Primers used for site-directed mutagenesis of bicistronic vectors.

**Supplementary Figure S10.**
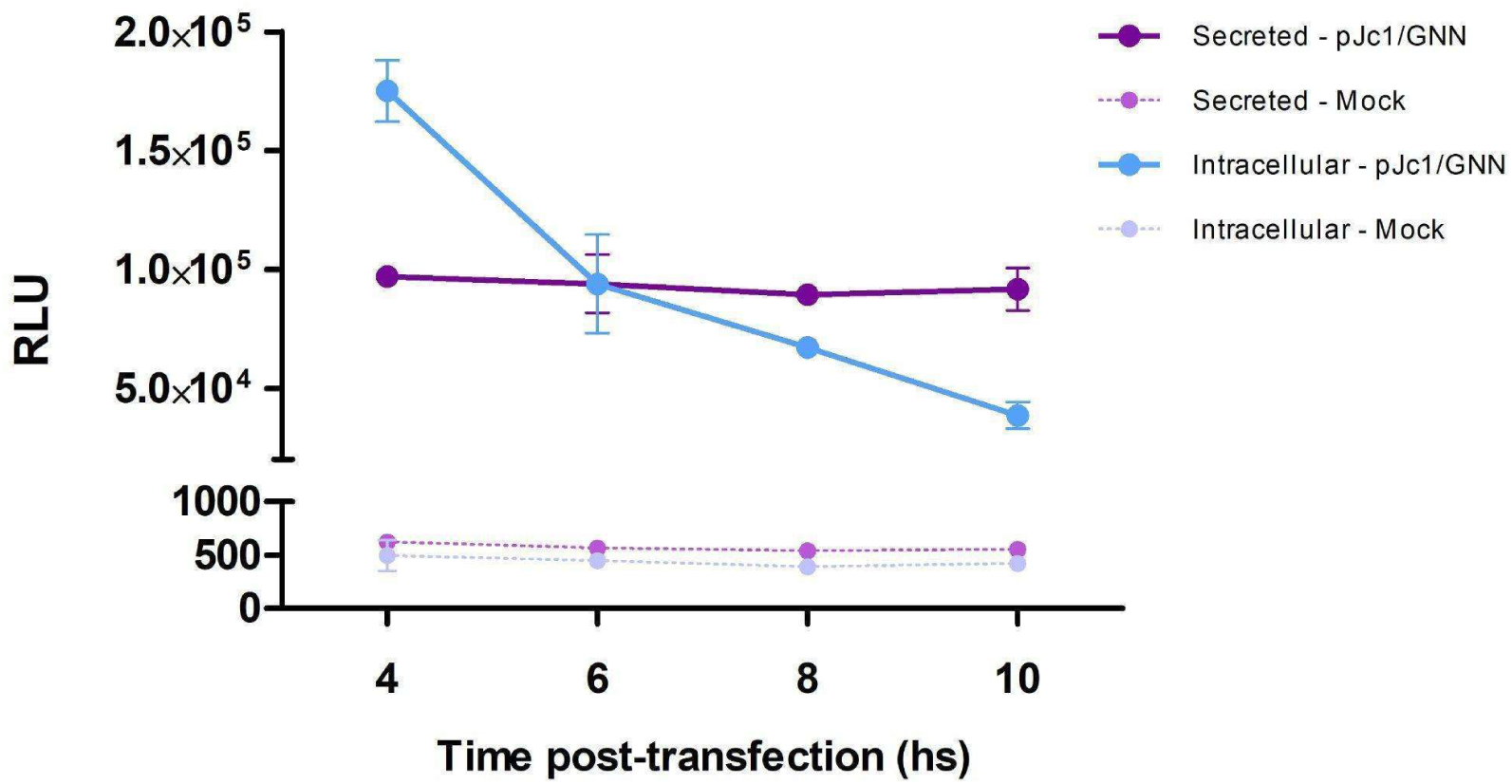
Detection of intra- and extracellular GLuc bioluminescence at different times post-transfection of the viral reporter RNA. Huh-7.5 cells were transfected with 100ng of viral reporter RNA (pJc1/GNN, coding for GLuc). At the indicated times, supernatants were collected and cells were lysed. The Renilla Luciferase Assay substrate was employed to quantify GLuc activity in each sample, with relative light units (RLU) being quantified with a FLUOstar Omega multimode plate reader (BMG Labtech). The results of three biological replicates are plotted. RLU: Relative Light Units.

